# A population-scale landscape of the subgingival microbiome reveals divergent routes to periodontal dysbiosis

**DOI:** 10.64898/2026.09.24.754177

**Authors:** Lu Li, Moemen Hussein, Karen L. Falkner, Robert J. Genco, Yijun Sun, Jean Wactawski-Wende, Michael J. Buck, Haralampos Hatzikirou, Patricia I. Diaz

**Author notes:** Corresponding Author: Patricia I. Diaz. Equal contribution (Lu Li, Moemen Hussein). Author is deceased (Robert J. Genco).

## Abstract

Periodontitis is an archetypical mucosal inflammatory disease in which microbiome dysbiosis at the tooth-epithelial interface interacts with host genetic and behavioral risk factors to drive immune-mediated tissue destruction. Although subgingival microbiome compositional shifts are thought to parallel disease severity, microbiome variation at the population-level and its relationship to periodontal clinical phenotypes and disease-modifying factors remain poorly defined. Here, we use unsupervised manifold learning to map the compositional landscape of the subgingival microbiome in 1,355 adults spanning periodontal health to severe periodontitis. We identified eight latent microbiome states organized along a branching continuum from eubiosis to dysbiosis. An intermediate microbial configuration marked ecological destabilization and bifurcation into two distinct periodontitis-associated dysbiotic trajectories, distinguished by links to gingival inflammation and smoking. Although the microbiome trajectories broadly tracked periodontal destruction, a minority of individuals showed discordant microbiome-clinical phenotypes, with some individuals with periodontitis retaining otherwise eubiotic microbiomes enriched for low-abundance pathobionts, while some cases of health or mild disease had highly dysbiotic communities, suggesting distinct host susceptibility. Together, these findings define a population-scale ecological landscape of the subgingival microbiome, reveal divergent trajectories to periodontal dysbiosis, and highlight heterogeneity in the relationship between microbial community structure and clinical disease expression.

## Background

Periodontitis is a bacterially induced chronic inflammatory disease that leads to progressive destruction of the tooth-supporting alveolar bone and soft tissues. Landmark natural history studies show that periodontal deterioration does not occur uniformly across individuals. Even in the absence of professional dental cleaning or deliberate preventive efforts, some individuals maintain long-term periodontal health, whereas others experience progressive tissue breakdown at variable and often episodic rates [1, 2]. This inter-individual heterogeneity likely reflects a complex interplay among the subgingival microbiome, host genetic and immune factors and cumulative environmental exposures [3].

At the population level, cross-sectional epidemiological data reveal a broad distribution of periodontal disease severity. In the United States, approximately 42.2% of adults are affected by periodontitis, yet only 7.8% present with severe disease [4]. This disparity indicates that most affected individuals reside within mild-to-moderate states rather than at the extreme end of tissue destruction. Case-control microbiological studies have established that periodontitis is accompanied by reproducible taxonomic shifts in the subgingival microbiome characterized by expansion of anaerobic and proteolytic taxa, including members of the so-called red complex such as *Porphyromonas gingivalis*, *Tannerella forsythia*, and *Treponema denticola*, and other inflammation-associated pathobionts [5–7]. However, despite detailed characterization of disease-associated taxa, how subgingival microbial communities are organized across the full spectrum of periodontal phenotypes remains poorly understood. In particular, it is unclear whether increasing disease severity is accompanied by a continuous ecological transition from eubiosis to dysbiosis, whether distinct dysbiotic configurations exist, and whether microbiome and clinical phenotypes are invariably concordant.

The relationship between microbiome dysbiosis and clinical disease is further complicated by factors that modify susceptibility to periodontitis. Cigarette smoking and diabetes mellitus profoundly influence periodontal disease risk and severity through effects on host inflammatory responses and the local microbial environment [8, 9]. Additional factors such as cardiovascular conditions, metabolic dysregulation and obesity, alcohol consumption, and low physical activity have also been associated with susceptibility to periodontitis [10–14]. Importantly, these modifiers may also reshape the subgingival microbiome by altering host immune surveillance and oxygen and nutrient availability within the periodontal niche [15–17]. Thus, variation in host factors and environmental exposures could contribute to heterogeneity in the relationship between microbial community structure and clinical disease expression.

Two major limitations constrain current understanding of subgingival microbiome organization in periodontitis at the population-level. First, most microbiome studies rely on case–control comparisons between periodontal health and advanced disease, emphasizing the extremes of the clinical spectrum while underrepresenting the much larger population with intermediate phenotypes. Such designs may obscure transitional microbiome configurations and preclude assessment of whether microbiome dysbiosis and clinical disease severity vary in parallel across the periodontal disease spectrum. Second, individuals with systemic comorbidities and major disease-modifying exposures are frequently excluded from these comparative studies, limiting insight into how real-world host variability shapes microbiome structure. Consequently, it remains unclear how microbial communities are distributed across the spectrum of periodontitis phenotypes and whether different clinical and host contexts are associated with distinct paths from eubiosis to dysbiosis.

Analytical approaches traditionally used to characterize microbiome variation impose additional constraints. Conventional microbiome analyses typically rely on beta-diversity measures coupled with linear ordination techniques to summarize compositional variation. While informative, these approaches assume that biological heterogeneity can be adequately captured through global distances or orthogonal projections. Microbiome data, however, are high-dimensional, sparse, and shaped by nonlinear ecological processes. When community composition evolves along continuous or branching trajectories rather than as discrete clusters, linear projections can collapse intermediate configurations and obscure local geometric relationships that capture gradual ecological transitions. Analytical frameworks that preserve local neighborhood structure may be required to recover the latent topological organization of microbial communities across a population.

Nonlinear manifold learning provides an alternative framework for resolving such structure by representing high-dimensional observations within a lower-dimensional space while preserving local topology and relationships among neighboring community configurations. Applied to large populations, manifold-based approaches can reveal latent patterns of microbiome variation and identify continuous trajectories, branching structures, and intermediate states that may not be evident from conventional ordination or discrete clustering. This framework therefore offers an opportunity to map the population-level ecological landscape of the subgingival microbiome across periodontal phenotypes and disease-modifying exposures. Similar manifold-based strategies have been used to characterize gut microbial configurations in Crohn’s disease reconstructing dysbiotic trajectories that track disease severity and periods of disease exacerbation and remission [18].

In this study, we leverage a comprehensive population-based cohort spanning the full clinical spectrum of periodontal health and disease in adults. By integrating microbial community data with a machine-learning pipeline incorporating unsupervised feature selection, nonlinear manifold learning, and graph-based clustering, we reconstruct a high-order ecological landscape of the subgingival microbiome defining its population-level topology. This approach revealed microbiome configurations embedded within continuous and branching trajectories from eubiosis to dysbiosis and quantified the association between host modifiers and these transitions. By moving beyond traditional case–control dichotomies and linear ordination approaches, our work provides a population-scale view of subgingival microbiome organization and its relationship to periodontal clinical phenotypes and host factors.

## Results

### Broad associations of microbiome and periodontal health status are detected by traditional diversity metrics, but fine-scale variation remains unresolved

To characterize microbiome variation in relation to periodontitis phenotypes and host factors, we integrated comprehensive clinical and host information collected from 1,355 adults with species-level microbiome composition, as determined by 16S rRNA gene amplicon sequencing (Fig. 1a). Subject enrollment was independent of periodontal health status; therefore, the cohort included subjects with the full spectrum of periodontal phenotypes in adults spanning from health to severe periodontitis. Cohort characteristics stratified by periodontal health status according to the Centers for Disease Control and Prevention and American Academy of Periodontology (CDC/AAP) surveillance definition [19] are shown in Supplementary Table S1. The prevalence of none/mild, moderate, and severe periodontitis was 23.0%, 49.8%, and 27.2%, respectively. The severe periodontitis group had a significantly higher proportion of males, current smokers, subjects with a history of myocardial infarction, subjects with self-reported high cholesterol, and subjects taking anti-hypertensive and statin medications. Subjects with severe periodontitis were also less likely to visit the dentist annually, and had, on average, fewer years of education and older age than the less severe groups. In addition, we utilized an alternative disease classification based on maximum clinical attachment loss in line with the AAP case definitions (2017 World Workshop on the Classification of Periodontal and Peri-Implant Diseases and Conditions [20]) to perform an additional cohort stratification (Supplementary Table S2), and utilized a custom classification that incorporates gingival bleeding to distinguish periodontal health from gingivitis (see Methods) (Supplementary Table S3). These supplementary classification schemes revealed comparable distributions of periodontitis severity and associations with host characteristics.

**Figure 1.**
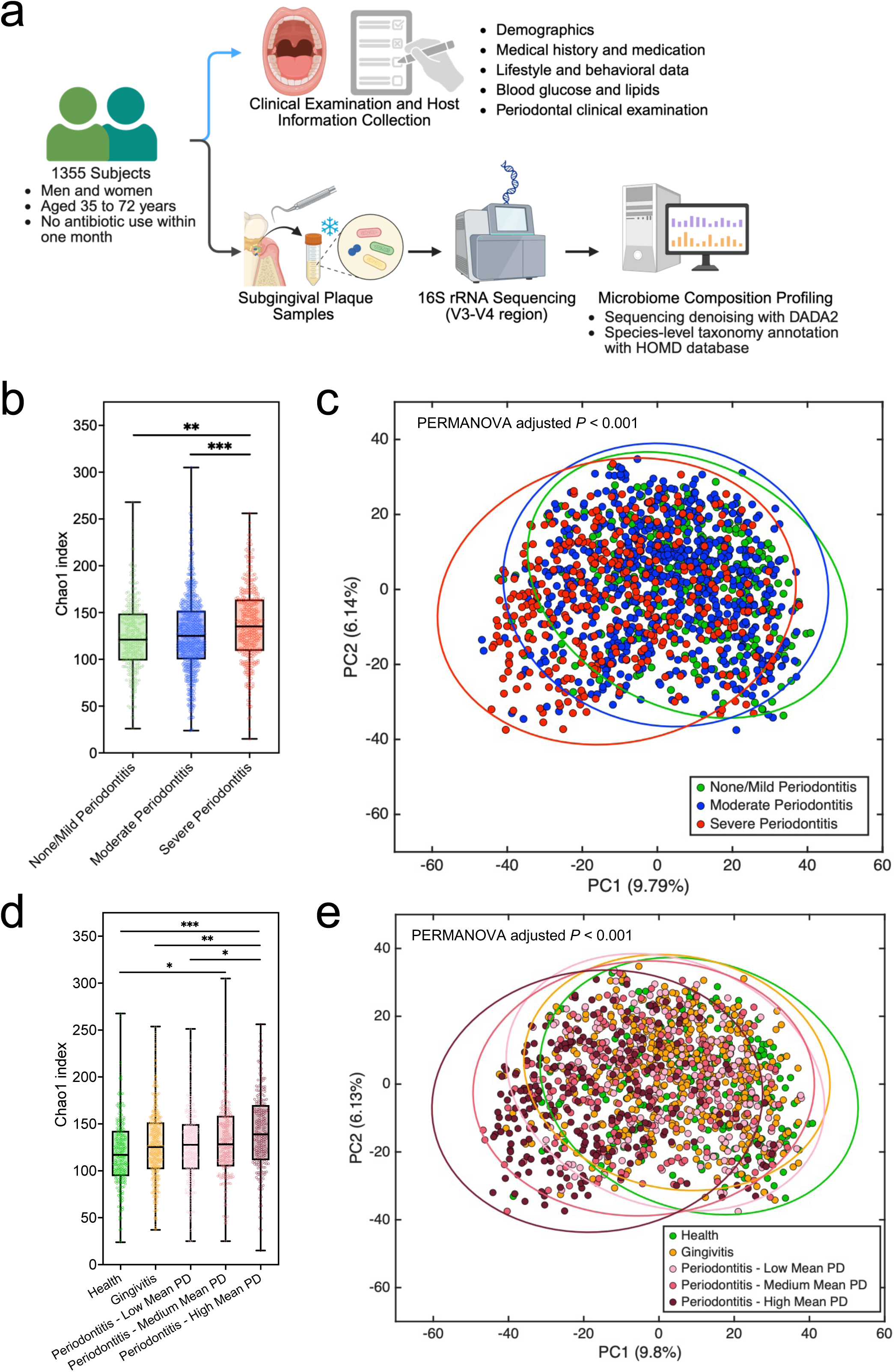
Species-level alpha and beta diversity show highly overlapping subgingival microbiome communities across periodontal health states. (a) Overview of the study design. Clinical and host information were collected from 1,355 subjects, and subgingival plaque samples were obtained for 16S rRNA gene sequencing to generate species-level taxonomic profiles. (b) Species-level alpha diversity according to CDC/AAP periodontitis groups. (c) Principal components analysis (PCA) of beta diversity based on Euclidean distances among CDC/AAP groups. (d) Alpha diversity of subjects categorized according to custom periodontal health criteria including health, gingivitis and periodontitis. (e) PCA of beta diversity based on Euclidean distances among custom-defined periodontal groups. Statistical significance for alpha diversity was assessed using ANCOVA with Bonferroni post-hoc correction, and for beta diversity using conditioned PERMANOVA. Models were adjusted for age, sex, years of education, history of myocardial infarction, current smoking status, lifetime smoking exposure in pack-years, lifetime alcohol consumption, annual dental visits, and daily flossing. * = *P* < 0.05, ** = *P* < 0.01, *** = *P* < 0.001.

To assess microbiome variation according to periodontal health status, we first employed alpha-and beta-diversity metrics. Subjects with severe periodontitis exhibited significantly higher alpha diversity (within-sample richness) compared to healthy individuals or those with milder disease (Figs. 1b and d, Supplementary Fig. S1a), consistent with previous findings [6, 21–23], although a wide range of alpha-diversity values was observed within each group. Beta-diversity (between-sample differences in community composition) analysis showed substantial overlap in global microbiome composition among groups (Figs. 1c and e; Supplementary Fig. S1b), although subjects with severe periodontitis exhibited partial separation from those classified as healthy, also in agreement with prior studies (Supplementary Figs. S2) [6, 7, 23]. These results indicate that although traditional diversity metrics capture associations of the subgingival microbiome and periodontal health status, particularly when considering the healthier and most severely diseased states, they show for the most part a rather homogeneous data distribution with significant overlap among periodontal health categories.

### Graph-based manifold learning reveals a branching eubiosis-to-dysbiosis continuum in the subgingival microbiome

To evaluate whether finer-scale and biologically relevant subgingival microbiome heterogeneity could be revealed using an alternative analytical approach, we applied a machine-learning–based framework, similar to that used in our previous work [18] but implemented here in an unsupervised manner. This framework operates in an agnostic manner relying solely on microbiome composition to reveal the hidden structure embedded in the data and does not incorporate host characteristics during modeling. Our analytical framework comprised three conceptual steps (Fig. 2a). First, unsupervised feature selection was performed using a structure-aware algorithm (iDetect [24]) to retain microbial taxa that captured informative variation and local neighborhood relationships among samples, without relying on a priori clinical labels. Second, the high-level organization of samples, based on centered log_2_-ratio (CLR)-transformed abundances of the selected species, was reconstructed using nonlinear manifold learning and reversed graph embedding (DDRTree [25]). This step inferred a three-dimensional principal-tree that reconstructed the latent topological organization of microbiome communities across the population, capturing continuous transitions and branching trajectories while preserving local neighborhood relationships among samples. This approach was motivated by initial beta-diversity analyses showing a graded continuum of community configurations across periodontal phenotypes, rather than discrete, well-separated clusters. Third, to facilitate the biological and clinical interpretation of this continuous landscape, spectral clustering was applied directly to the low-dimensional manifold embedding, partitioning samples along the inferred structure into discrete, topologically cohesive microbiome configurations representing interpretable states along the microbiome continuum (see Methods for details).

**Figure 2.**
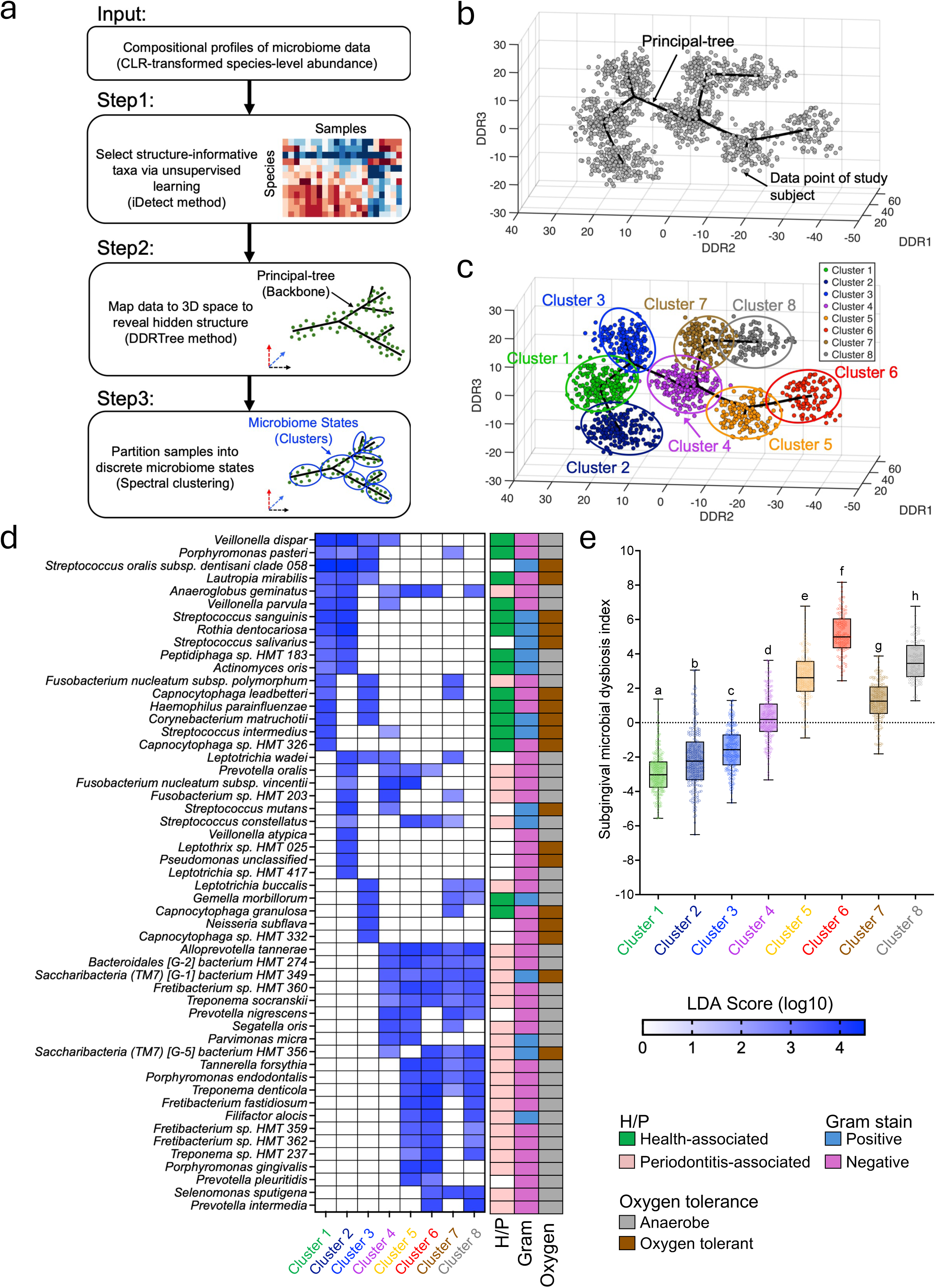
Graph-based manifold learning framework uncovers latent microbiome states arranged along a branching eubiosis-to-dysbiosis continuum. (a) Schematic overview of the analytical workflow. Structure-aware unsupervised feature selection using iDetect was applied to identify informative taxa. CLR-transformed abundances of selected species were then projected into a three-dimensional manifold using DDRTree to reconstruct a latent principal-tree summarizing microbiome variation, followed by spectral clustering to identify discrete microbiome states. (b) Three-dimensional principal-tree reconstruction. The manifold was inferred using DDRTree based on the CLR-transformed abundances of 599 iDetect-selected species. Each point represents an individual subject, and the solid line denotes the principal-tree capturing the primary structure of microbiome variation. (c) Eight discrete clusters were defined using spectral clustering based on geodesic distances along the inferred principal tree. (d) Heatmap of cluster-specific microbial signatures. Differentially enriched species were identified using LEfSe by comparing each cluster against all remaining clusters. Displayed species had a Linear Discriminant Analysis (LDA) score ≥3.5 in at least one cluster. (e) Box plots showing the distribution of the Subgingival Microbial Dysbiosis Index (SMDI) across the eight clusters. Statistical significance was assessed using one-way ANOVA with Bonferroni-adjusted post-hoc pairwise comparisons. All pairwise comparisons were significant (*P* < 0.05), as indicated by the distinct letters above the boxes.

Applying this framework revealed a trajectory with multiple branching points, indicative of heterogeneous microbiome configurations within the population (Fig. 2b). Because the cohort includes a substantial number of individuals who had recently survived a myocardial infarction (MI), and the influence of MI or its associated comorbidities and treatments on the subgingival microbiome is incompletely understood, we evaluated whether MI status affected the inferred structure. When stratified by recent MI history, participants were broadly distributed across the inferred landscape, with no distinct MI-associated branch or cluster (Supplementary Fig. S3a). Repeating the analysis after restricting the cohort to participants without a history of MI recovered a similar broad topology, comprising a left-to-right principal tree with two major branches (Supplementary Fig. S3b). This finding indicates that the major branching architecture was not an artifact of including participants with a recent MI history and suggests that it may generalize beyond MI-enriched cohorts.

Spectral clustering analysis identified eight microbiome states (clusters), each corresponding to a specific region of the learned structure providing a framework for evaluating and interpreting community-level microbiome variation across the population (Fig. 2c). To explore the distinguishing features of each microbiome state, we performed linear discriminant analysis effect size (LEfSe) analysis defining taxa differentially enriched in each cluster in comparison to the remainder of the cohort (Fig. 2d, Supplementary Table S4). Clusters 1, 2 and 3 were enriched for several commensals previously associated with periodontal health. In contrast, a variety of species with known association with periodontitis were enriched in Clusters 5 to 8. Cluster 4, located at the mid-branching point in the tree structure, exhibited a mixed signature, with enrichment of health-associated commensals alongside multiple periodontitis-associated pathobionts. Notably, LEfSe analysis also revealed species differentially enriched in each of the two right branches. For instance, the periodontitis-associated species *Porphyromonas gingivalis* was enriched exclusively in the lower-branch clusters (Clusters 5 and 6), whereas species, such as *Leptotrichia buccalis* and *Selenomonas sputigena*, previously linked to gingivitis [6, 26], were enriched in both upper branch clusters (Cluster 7 and 8) and in the terminal point of the lower branch (Cluster 6), but did not appear enriched in the lower branch Cluster 5.

Further examination of the most frequently detected and abundant species in each cluster (Supplementary Fig. S4) showed Cluster 1–3 were indeed dominated by commensal symbionts, whereas Clusters 5–8 showed predominance of Gram-negative anaerobes commonly linked to periodontitis [6], and Cluster 4 exhibited a mixture of health-and disease-associated taxa. In addition, some species such as *Streptococcus oralis subsp. dentisani clade 058*, *Veillonella dispar*, and *Fusobacterium nucleatum subsp. vincentii* were consistently dominant across all states (Supplementary Fig. S4), supporting their proposed status as core members of the subgingival microbiome that are prevalent and abundant regardless of periodontal health [23, 26].

Since the preceding analyses suggested a eubiosis-to-dysbiosis progression, from left to right, along the inferred tree structure, we quantified these shifts across clusters using the subgingival microbial dysbiosis index (SMDI), a previously defined score based on the mean CLR abundance of periodontitis-associated species minus that of health-associated species, with higher values indicating greater dysbiosis [27]. SMDI values differed significantly across clusters (Fig. 2e). The left Cluster 1 showed the lowest values, with Clusters 2 and 3 showing modest increases. SMDI increased progressively toward both the right endpoints of the tree structure, with Cluster 6 in the lower branch and Cluster 8 in the upper branch showing the highest scores. This gradient provides additional evidence that the inferred trajectory captures a continuous left-to-right shift in microbiome composition from eubiosis to dysbiosis along the X-axis (DDR2 in Fig. 2c).

Altogether, these findings suggest the identified low-dimensional tree structure captured biologically relevant, continuous microbiome states, with Cluster 4 acting as a branching point from which two ecological routes leading to distinct dysbiotic communities emerge.

### Microbiome eubiosis to dysbiosis continuous states correlate with periodontal disease severity and are associated with distinct host factors

To investigate the clinical relevance of the identified microbiome states, we evaluated periodontal clinical parameters and host factors that differed across clusters using univariate analysis. Fig. 3a summarizes the average periodontal clinical measures and host variables that differed significantly. The microbiome configurations along the inferred left-to-right trajectory, corresponding to a shift from eubiosis to dysbiosis, also aligned with a clear gradient of periodontal disease severity. Clinical parameters indicating periodontitis were lowest in Cluster 1, followed by Cluster 2, and 3, consistent with SMDI gradients (Figs. 3a and 2e). Indicators of periodontitis increased significantly in Cluster 4, rising further along both right branches, reaching higher values in Cluster 8 on the upper branch and the further increasing in severity in Cluster 6 on the lower branch, in full agreement with SMDI values (Figs. 2e, 3a-c and Supplementary Fig. S5).

**Figure 3.**
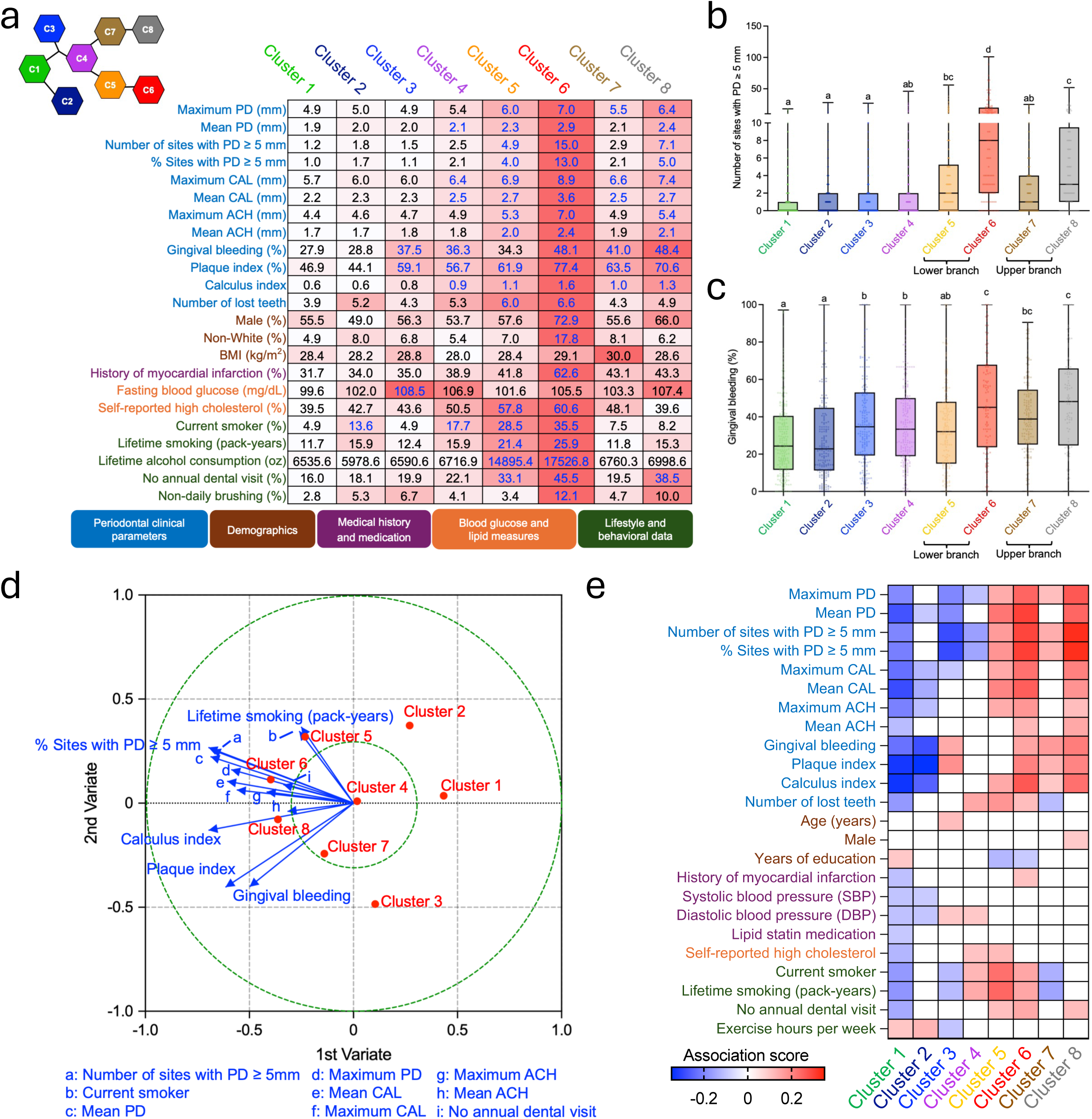
Association of microbiome clusters with clinical periodontal states and with demographic, medical and behavioral subject characteristics. (a) Heatmap showing average values of periodontal clinical parameters, demographic characteristics, medical history and medication variables, and lifestyle and behavioral factors that differed significantly across clusters. For continuous variables, significance was assessed using one-way ANOVA; for categorical variables, significance was assessed using chi-squared tests. Bonferroni correction was applied to account for multiple testing. Clusters significantly different from Cluster 1 are highlighted in blue. Cell shading indicates the relative magnitude of each variable across clusters, ranging from white (lower) to red (higher). (b) Box plots showing the distribution of number of sites with PD ≥5mm across clusters. (c) Box plots showing the distribution of gingival bleeding across clusters. (d) Regularized canonical correlation analysis (rCCA) of associations between microbiome clusters and host factors. The biplot shows correlations of microbiome clusters and host factors with the first two canonical variates. Microbiome clusters are shown as red points, and host factors are shown as blue arrows. Only host factors with a distance greater than 0.3 from the origin are shown. (e) Heatmap showing rCCA-derived association scores between host factors and microbiome clusters. Only scores with an absolute value ≥0.09 are displayed.

Beyond periodontal parameters, several lifestyle, demographic, and medical factors also varied significantly across clusters (Fig. 3a; Supplementary Figs. S6). Cluster 6, the most dysbiotic state and the cluster with the most severe periodontitis, was associated with higher prevalence of myocardial infarction and self-reported high cholesterol, greater smoking and alcohol exposure, limited dental care and oral hygiene, and higher proportions of males, and non-white individuals, all factors previously associated with periodontitis [4, 28–32]. Fasting blood glucose and BMI, two parameters previously linked to periodontitis [33–35], showed limited associations with the microbiome clusters (Fig. 3a; Supplementary Figs. S6). Fasting blood glucose was lowest in Cluster 1, the most eubiotic state, and highest in Cluster 3 with other clusters showing limited variation. BMI showed limited variation across clusters, suggesting a weak association with the identified microbiome states.

### Gingival bleeding and smoking are the main host factors associated with two distinct dysbiotic trajectories

Examination of the two branches emerging from Cluster 4 revealed distinct associations with host factors (Fig. 3a-c and Fig. S6). Gingival bleeding was significantly higher in both the moderately and highly dysbiotic clusters (Clusters 7 and 8) of the upper branch when compared to the more eubiotic clusters 1 and 2, which showed low bleeding. In contrast, along the lower branch, bleeding was significantly increased, relative to Clusters 1 and 2, only at the terminal cluster (Cluster 6), whereas the moderately dysbiotic Cluster 5 did not differ significantly from the eubiotic Clusters 1 and 2 in terms of bleeding (Fig. 3a–c). Gingival bleeding is a sign of local inflammation that commonly accompanies periodontitis, but can also occur irrespective of the presence of irreversible periodontal damage (ie. in gingivitis). Accordingly, gingival bleeding was also increased in the eubiotic Cluster 3 compared to Clusters 1 and 2, altogether suggesting that the Y axis of the tree structure (DDR3 in Fig. 2c) captured microbiome associations with bleeding.

In contrast, smoking-related variables were preferentially associated with the lower branch, particularly with Clusters 5 and 6, while the upper branch clusters had similar levels of smokers as the eubiotic clusters. Together, these findings suggest that the two microbiome routes leading to two distinct terminal dysbiotic states associated with severe periodontitis differ in their association with gingival inflammation and smoking habit. Moreover, current smoking was the only host parameter that differed between Clusters 1 and 2, suggesting this environmental exposure may also modify eubiotic microbiome communities.

To assess multivariate relationships between host factors and microbiome states, we performed regularized canonical correlation analysis (rCCA). The rCCA biplot showed that periodontal clinical measures were the dominant contributors to variation across microbiome states, with smoking-related variables and gingival bleeding also loading strongly along the canonical variates (Fig. 3d). Both current smoking and lifetime smoking exposure (pack-years) loaded predominantly alongside the lower-branch clusters (Clusters 5 and 6), particularly near Cluster 5, whereas gingival bleeding loaded alongside the upper branch clusters (Clusters 7 and 8), consistent with the univariate patterns. The association heatmap further illustrated how individual host factors related to each microbiome state, with current smoking and pack-years showing the strongest associations with lower-branch clusters and gingival bleeding mapping primarily to upper-branch clusters (Fig. 3e). These multivariate results strengthen the evidence that the two trajectory branches represent distinct dysbiotic pathways influenced by different host factors.

To further investigate ecological differences between the two trajectory branches, which differed in their associations with gingival bleeding and smoking, we examined species distinguishing their moderately dysbiotic states represented by Cluster 7 in the upper branch and Cluster 5 in the lower branch. LEfSe analysis revealed clear branch-specific enrichments consistent with the host-associated patterns described above (Fig. 4a). Cluster 7 was enriched for taxa previously linked to gingivitis, including *Fusobacterium nucleatum subsp. polymorphum*, *Selenomonas sputigena*, and several *Leptotrichia spp.*, such as *L. buccalis* [6, 26]. By contrast, Cluster 5 was enriched for classic periodontitis-associated species, consistent with a more severely dysbiotic configuration (Fig. 2e), and for taxa previously associated with smoking, such as *Fusobacterium nucleatum subsp. vincentii* [36, 37]. Although fewer species distinguished the two branches in their terminal states (Clusters 6 and 8), the same ecological contrast remained evident (Fig. 4b). The upper branch continued to be characterized by taxa associated with gingivitis, whereas the lower branch remained enriched for classic periodontal pathobionts and species associated with smoking. Notably, the dominant discriminative signal in the lower branch shifted from *Fusobacterium nucleatum subsp. vincentii* at the moderately dysbiotic state (Cluster 5) to *Porphyromonas gingivalis* at the terminal state (Cluster 6), suggesting a change in ecological dominance from a bridging species to a more virulent pathogen across the inferred dysbiosis gradient.

**Figure 4.**
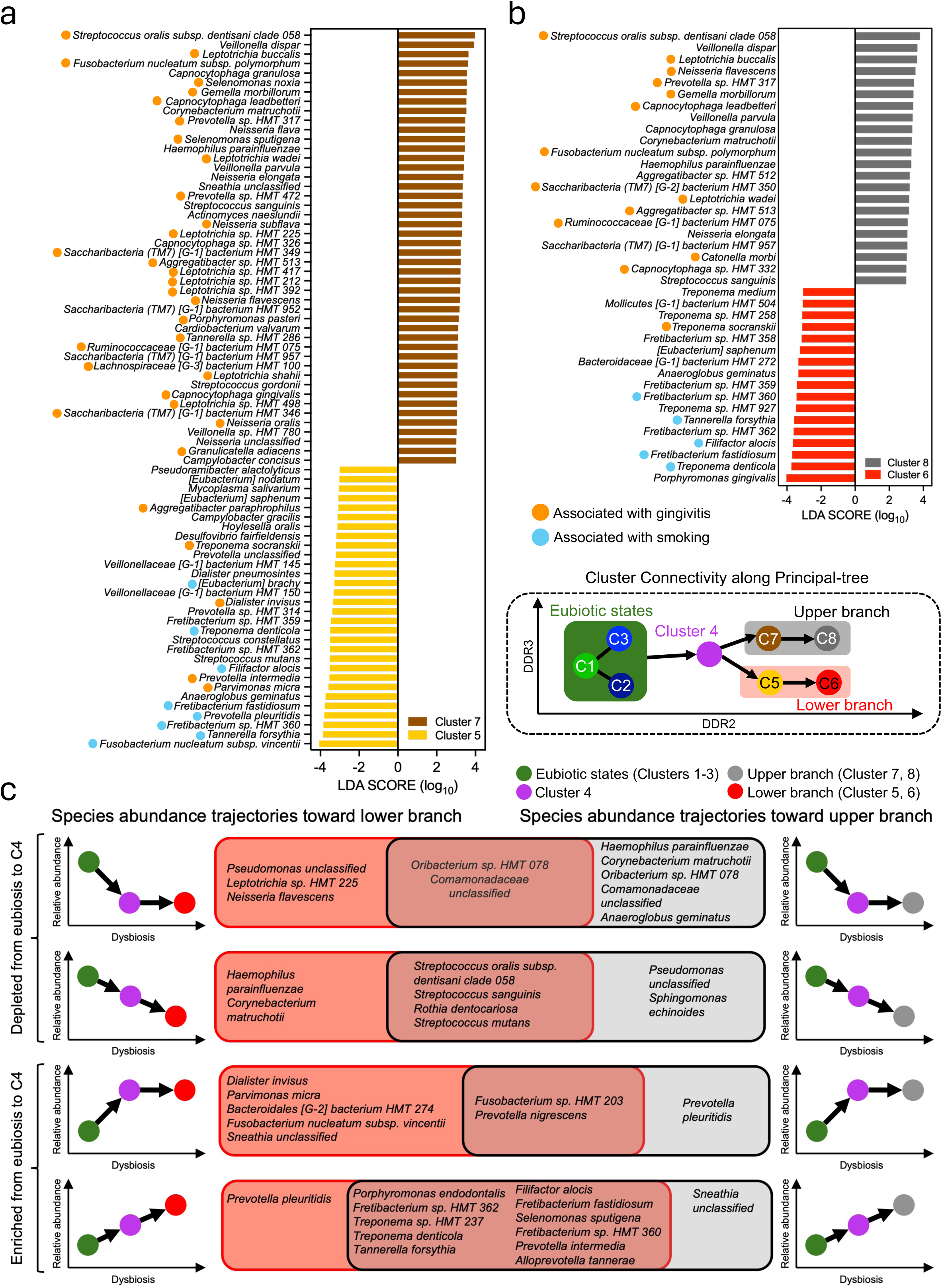
Species abundances along the principal-tree highlight distinct ecological trajectories from health toward dysbiosis. (a) LEfSe analysis comparing the moderately dysbiotic lower-branch (Cluster 5) with the moderately dysbiotic upper-branch (Cluster 7). (b) LEfSe analysis comparing the highly dysbiotic lower (Cluster 6) and upper (Cluster 8) branches. Species previously associated with gingivitis are indicated by orange dots, and smoking-associated species identified by rCCA analysis (Supplementary Fig. S7) are annotated in blue. (c) Differential species identified by LEfSe as enriched or depleted when comparing eubiotic states to the intermediate Cluster 4, grouped according to whether their trajectories subsequently remained at similar levels or became further enriched or depleted in the upper or lower dysbiotic branches.

### Species-specific associations confirm periodontitis as the greatest influence in the subgingival microbiome, with gingival bleeding and smoking as additional modifying factors

To better understand associations of individual species with host metadata, independent of the microbiome community context, we applied rCCA to examine associations between individual species abundances and host factors (Supplementary Fig. S7). This analysis identified species known from previous studies [6, 23, 26] to have strong positive or negative associations with periodontitis. We also identified a set of species highly associated with gingival bleeding and plaque index but not associated with periodontitis showing that species associations with the destructive inflammatory process of periodontitis and associations with the reversible inflammatory process of gingivitis are distinct. This analysis also showed species negatively and positively associated with smoking, assessed by current smoking status and cumulative lifetime exposure, including commensals inversely associated with periodontitis and periodontitis-associated pathobionts, respectively. Most of the species positively-associated with smoking were also significantly enriched in the lower branch of the inferred learning tree structure (Fig. 4a), relative to the upper branch, reinforcing smoking as a key ecological correlate of this trajectory. Taken together, species-level analyses showed that periodontal status was most strongly associated with subgingival microbiome composition, with additional associations observed for gingival bleeding and smoking, whereas the remaining measured host variables exert minimal effects.

### Cluster 4 is a branching point where health-associated taxa decline, while pathobionts expand

We next evaluated the ecological position of Cluster 4, which represents the bifurcation point leading to the two dysbiotic branches. Several health-associated taxa, such as *Haemophilus parainfluenzae*, *Corynebacterium matruchotii* and *Streptococcus sanguinis* were depleted in Cluster 4 compared with the eubiotic states, with the abundance of these species either stabilizing or further declining in the more dysbiotic branches (Fig. 4c). This pattern indicates Cluster 4 marks a loss of community structure characteristic of periodontal health. Furthermore, a subset of species from the genera *Fusobacterium* and *Prevotella*, peaked in Cluster 4 and thereafter remained elevated in both dysbiotic branches (*Fusobacterium* sp. HMT 203, *Prevotella nigrescens* and *Prevotella intermedia*; Fig. 4c), or declined in the dysbiotic states but still remaining above eubiotic levels (*F. nucleatum* subsp. *animalis* and *Prevotella oris*; Supplementary Fig. S8). In parallel, red complex taxa, including *Treponema denticola* and *Tannerella forsythia*, and additional periodontitis-associated pathobionts, including *Filifactor alocis*, *Porphyromonas endodontalis*, *Prevotella intermedia*, *Fretibacterium fastidiosum*, and other *Fretibacterium spp.*, became enriched in Cluster 4 and further increased along both dysbiotic branches (Fig. 4c). These coordinated patterns indicate that Cluster 4 marks an intermediate ecological state in which health-associated community structure begins to collapse, bridging taxa such as *Fusobacterium* spp. expand, and key pathobionts gain momentum before the community separates into distinct dysbiotic trajectories. This ecological configuration provides the foundation from which the upper and lower branches emerge.

### Low abundance pathobionts are associated with periodontitis in individuals with an otherwise health-like microbiome configuration

While we identified an eubiosis-to-dysbiosis continuum correlated with the transition from periodontal health to periodontitis, the clinical parameters of subjects within each cluster varied considerably (Supplementary Fig. S5), suggesting microbiome-clinical phenotype relationships are not always concordant. Examination of the correlations between microbiome dysbiosis index and clinical parameters showed that although most subjects had concordant clinical disease-microbiome dysbiosis relationships, as evidenced by positive and significant correlations between these parameters, a small proportion of individuals appear as outliers (Fig. 5a; Supplementary Fig. S9). That is, there are individuals with relatively low dysbiosis despite substantial periodontal damage and subjects with highly dysbiotic communities and clinical parameters indicative of health or only mild disease. Evaluation of disease cases by cluster showed that although most subjects in the eubiotic clusters (1, 2 and 3) are classified as periodontally healthy or having only gingivitis, a minority of subjects in these clusters showed signs of periodontitis with a small proportion (3 to 9%) showing severe disease (Fig. 5b).

**Figure 5.**
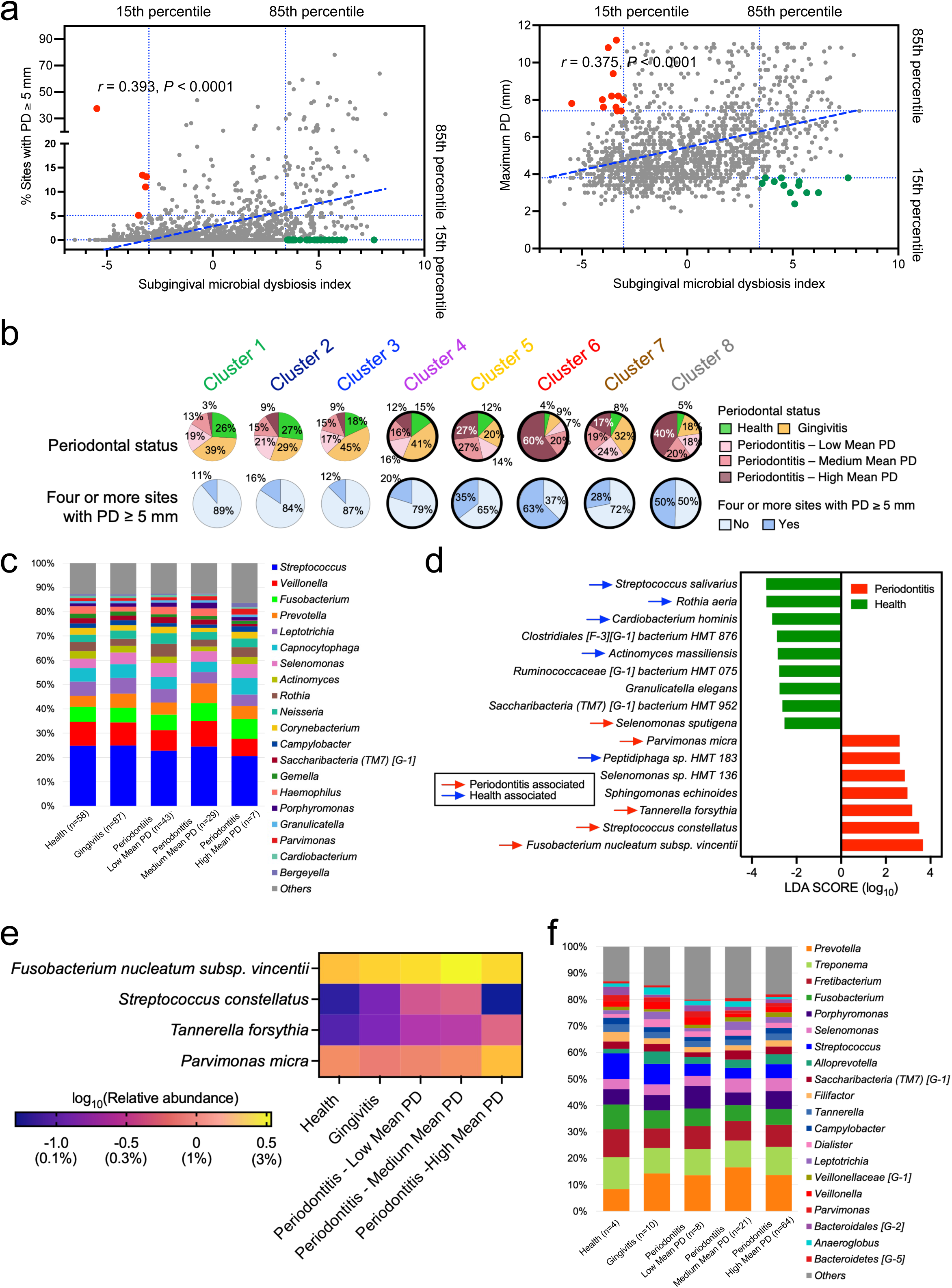
Eubiosis and dysbiosis are decoupled from periodontal clinical phenotypes in a minority of subjects. (a) Scatter plots show Pearson correlations between the subgingival microbial dysbiosis index (SMDI) and the percentage of sites with PD ≥5 mm or maximum PD. Green points indicate individuals with high dysbiosis but low signs of clinical disease, defined as SMDI above the 85th percentile and the corresponding clinical measure below the 15th percentile. Red points indicate individuals with low dysbiosis but high clinical disease, defined as SMDI below the 15th percentile and the corresponding clinical measure above the 85th percentile. (b) Pie charts show the distribution of clinical groups across the eight microbiome clusters based on the custom periodontal classification and presence of ≥ 4 sites with PD ≥ 5 mm. Statistical significance was assessed using chi-squared tests with Bonferroni correction for multiple comparisons. Clusters significantly different from Cluster 1 are shown in bold. (c) Top 20 most abundant genera in Cluster 1 subjects classified as health, gingivitis, and periodontitis showing similarity in community composition. (d) LEfSe analysis comparing health and periodontitis subjects within Cluster 1. Species previously associated with health and periodontitis are indicated by blue and red arrows, respectively. (e) Heatmap showing relative abundance of pathobionts differentially enriched in Cluster 1 subjects with periodontitis compared to periodontal healthy subjects. Enriched species were identified by LEfSe. (f) Relative abundance of the 20 most abundant genera within Cluster 6 across individuals with health, gingivitis, and periodontitis.

To further explore why some subjects with an eubiotic microbiome presented with severe periodontitis, we focused on Cluster 1, the most eubiotic community configuration. Detailed inspection of the clinical periodontal measures in the 3% of Cluster 1 subjects with severe periodontitis (7 subjects), confirmed they exhibited severe periodontal destruction (Supplementary Table S5). To evaluate whether all communities in Cluster 1 retained an eubiotic configuration, we visualized the most abundant genera across subjects in this cluster. Even among individuals with severe periodontitis, the dominant genera indicated a community structure similar to subjects with periodontal health (Fig. 5c, Supplementary Fig. 10). To assess whether subtle taxonomic differences were present in subjects with periodontitis, we performed LEfSe comparing the communities of Cluster 1 subjects with periodontitis versus those periodontally healthy (Fig. 5d). This analysis revealed enrichment of the periodontitis-associated pathobionts *Tanerella forsythia* and *Parvimonas micra* in diseased individuals, albeit these taxa were present in relatively low abundance (Fig. 5e). Taken together, these findings suggest that periodontitis in subjects with an apparently eubiotic microbiome is associated with enrichment of low-abundance canonical pathobionts.

### A minority of individuals showed highly dysbiotic microbiomes despite limited periodontitis

Cluster 6, which represents the most dysbiotic state, contained a small subset of individuals (13%) who did not meet the periodontitis definitions and were classified as healthy or having gingivitis (Fig. 5b). The microbial communities of these subjects closely resembled those of subjects with periodontitis within the same cluster, with high proportions of known pathobionts (Fig. 5f; Supplementary Fig. S11a). Species characteristically enriched in the highly dysbiotic Cluster 6 when compared to other clusters (LEfSe, Supplementary Table S4) did not differ significantly in abundance across Cluster 6 subjects with health, gingivitis, and periodontitis confirming all microbiomes were equally dysbiotic (Supplementary Fig. S11b).

One explanation for the incongruent clinical phenotype-microbiome relationship in Cluster 6 subjects classified as healthy or with gingivitis may be that these subjects were previously diseased and subsequently treated, with their periodontal pockets resolving but not their dysbiosis. Recent evidence indeed shows that current periodontal therapies are not entirely effective at resolving dysbiosis [38–43]. Close examination of the clinical parameters of Cluster 6 subjects classified as currently healthy or having gingivitis indeed showed evidence, in some of these subjects, of loss of clinical attachment, often accompanied by tooth loss, findings consistent with a history of prior periodontitis (Supplementary Table S6). However, not all Cluster 6 subjects classified as healthy or with gingivitis showed a reduced periodontium, with some individuals exhibiting clinical attachment levels compatible with health without an apparent history of periodontitis and no tooth loss. These findings indicate that highly dysbiotic microbiome configurations can be observed in individuals without current periodontitis and, in some cases, without evidence of an apparent history of prior periodontal destruction. Thus, severe dysbiosis does not necessarily coincide with severe inflammation or periodontal destruction, despite previous suggestions that inflammation is critical to sustain periodontitis-associated dysbiosis [44].

## Discussion

In this population-based study, we applied an unsupervised machine-learning framework, utilizing topology-aware feature selection and nonlinear graph-based manifold learning, to reconstruct a population-level high-order subgingival microbiome landscape. Rather than separating into discrete health-and diseased-associated clusters, microbial communities were organized along a branching continuum comprising eight configurations. Although the present study is based on cross-sectional study samples, the inclusion of a large cohort enabled reconstruction of population-level variability and inference of potential ecological trajectories. Whether the inferred trajectories reflect longitudinal transitions of individual microbiome communities will require prospective validation. Nevertheless, the inferred population-level architecture extends current models of periodontitis as a polymicrobial dysbiotic condition driven by community-level imbalance and host-microbe interactions [45], by suggesting that dysbiosis associated with severe disease may arise through multiple ecological routes rather than a single linear progression, while also positioning cigarette smoking as a central modifier of microbiome composition.

Although moderate periodontitis was the predominant diagnostic category in our cohort, consistent with national epidemiological trends [4], the overall prevalence of periodontitis exceeded nationally representative estimates from the National Health and Nutrition Examination Survey (NHANES) [4]. This discrepancy is unlikely to reflect differences in periodontal assessment, because both the NHANES and our study used CDC/AAP surveillance definitions derived from full-mouth six-site periodontal examinations. Rather, the higher prevalence of periodontitis in our cohort may reflect its case-control design and enrichment for individuals with myocardial infarction, who comprised 39.4% of participants. Periodontitis and myocardial infarction share several risk factors, including age, smoking, sex, and socioeconomic factors, and epidemiological studies have reported associations between periodontal disease and cardiovascular disease [46–48]. Therefore, while the prevalence of moderate periodontitis is consistent with population-level patterns, the higher absolute disease burden should be interpreted in the context of the cohort’s medically enriched sampling frame. This context does not diminish the observed organization of the subgingival microbiome landscape, but it should be considered when interpreting the distribution of clinical categories across microbiome states.

A key feature of the reconstructed landscape is the identification of Cluster 4 as an intermediate ecological state positioned at the juncture where eubiosis shifts toward dysbiosis. This transitional state was characterized by depletion of health-associated taxa and expansion of organisms linked to early dysbiotic reorganization, particularly members of the genera *Fusobacterium* and *Prevotella*. This pattern is consistent with the classic subgingival microbial complexes model of Socransky [5], in which orange-complex organisms, including *Fusobacterium* and *Prevotella* species, precede and facilitate colonization by red-complex pathogens. In our data, this ecological sequence was reflected by expansion of these bridging taxa at Cluster 4, followed by increased abundance of red-complex taxa and other periodontitis-associated pathobionts along the dysbiotic branches. The contribution of *Fusobacterium* and *Prevotella* to dysbiotic transitions is likely both structural and metabolic. *Fusobacterium* spp. are well-recognized bridging organisms that promote co-aggregation between early and late colonizers, thereby supporting biofilm integration and spatial organization [49–51]. In parallel, *Fusobacterium* spp. may provide metabolic support for periodontal pathogens through production of carbon-dioxide, which is utilized by pathogens such as *P. gingivalis* [52], and by creating strictly-anaerobic niches where pathogens can thrive [52]. In addition, *Fusobacterium* spp. have been shown to utilize amino acids supplied by other commensals and create a polyamine-rich environment that accelerates the biofilm life cycle of *P. gingivalis* [53]. There is also evidence that *Prevotella* spp. are avid biofilm formers [54], serve as bridging species [55], degrade host glycoproteins thereby modifying the local nutrient environment [56] and produce fermentation products—including succinate, acetate, and formate—that contribute to metabolic interactions among neighboring community members [57]. Conversely, depletion of *Corynebacterium matruchotii*, a taxon implicated in the formation of hedgehog-like spatial structures in supragingival plaque, may reflect loss of health-associated biofilm architecture [58]. Together, these findings suggest that transition beyond Cluster 4 reflects both weakening of health-associated ecological stability and emergence of a community configuration permissive to anaerobic pathobiont expansion. From a translational perspective, taxa and ecological configurations characteristic of this intermediate state may help identify individuals at increased risk of progression before the emergence of fully dysbiotic terminal communities. They may also inform therapeutic strategies aimed at preserving health-associated eubiosis, that by limiting expansion of bridging taxa disrupt metabolic interactions that support pathobiont outgrowth.

Beyond this transitional state, the manifold bifurcated into two major dysbiotic trajectories both associated with periodontitis but under distinct host-related contexts. The lower branch was preferentially associated with smoking and was characterized by a more severe dysbiotic configuration, greater disease severity, and enrichment of *P. gingivalis* and other periodontitis-associated virulent pathobionts (Fig. 2d-e, Fig. 3b; Supplementary Fig. S6). The upper branch was likewise associated with periodontitis but was distinguished by a closer association with gingival bleeding and enrichment of taxa previously associated with gingivitis (Fig. 2c, Fig. 3a and b). These findings suggest that dysbiosis is not a single endpoint but can develop along alternative ecological routes shaped by different modifying pressures. The smoking-associated lower branch provides a possible example of how environmental exposure may modify both microbial ecology and clinical disease expression. Smoking can suppress overt gingival bleeding through nicotine-induced vascular alterations that attenuate clinical signs of inflammation [59–61]. In parallel, smoking may favor *P. gingivalis*-associated dysbiosis by modifying both the host environment and the bacterial phenotype. Cigarette smoke extract has been shown to alter *P. gingivalis* virulence-related gene expression, including genes involved in fimbrial biogenesis and capsular polysaccharide synthesis, induce outer membrane proteins such as RagA and RagB, while reducing the proinflammatory response elicited by *P. gingivalis* in innate immune cells [62]. Moreover, *P. gingivalis* can subvert neutrophil-mediated immune control by suppressing IL-8-dependent neutrophil recruitment and chemotaxis, potentially contributing to muted clinical signs of gingival inflammation despite periodontal destruction [63, 64]. Together, these mechanisms may help explain the lower-branch phenotype, in which the moderately dysbiotic state (Cluster 5) showed elevated periodontal destruction without a corresponding significant increase in gingival bleeding, whereas bleeding became more evident at the terminal dysbiotic state (Cluster 6), once in the presence of severe periodontal destruction. By contrast, the upper branch was characterized by increased gingival bleeding and enrichment of gingivitis-associated taxa, supporting a distinct dysbiotic route more closely linked to overt gingival inflammation in conjunction with periodontal destruction.

Another important finding of our study was the presence of discordant clinical-microbiome phenotypes. Notably, a subset of individuals within the most eubiotic microbiome state presented with clinical periodontitis, including severe forms of the disease. Although their overall community structure remained dominated by health-associated taxa, differential analysis detected enrichment of low-abundance pathobionts including *T. forsythia* and *P. micra*. This pattern suggests that clinical disease can arise in the absence of overt community collapse, as low-abundance organisms have been previously suggested capable of exerting disproportionate immunomodulatory effects that lead to periodontitis [65]. Disease presentation in the presence of an overall eubiotic community enriched for low abundance pathobionts may also reflect increased host susceptibility since periodontitis is a polygenic disease with 30 to 50% of disease expression estimated to be due to genetic predisposition, mostly involving genes associated with healing, immune responses to the microbial insult and maintenance of structural integrity of the periodontal tissues [66, 67]. Conversely, a small subset of individuals within the most dysbiotic state (Cluster 6) presented with relatively healthy or mildly affected tissues despite harboring pathogen-enriched communities comparable to those of severely diseased subjects. These findings contradict the concept that inflammation is the critical factor that fuels dysbiosis, as highly dysbiotic communities were evident in individuals with non-inflammed tissues. These findings also suggest that individuals with periodontal health but highly dysbiotic communities may be tolerant to the challenge imposed by pathobionts. Alternatively, the dysbiotic microbiomes in these subjects may reflect an early ecological transition preceding clinically detectable destruction. Longitudinal studies are needed to distinguish these possibilities. Overall, our observations suggest that microbial burden alone is insufficient to determine clinical outcome and support the concept that host susceptibility modulates whether a dysbiotic community translates into overt periodontal destruction [68–70]. Furthermore, we observed that individuals with mild or moderate periodontitis did not exhibit a single severity-specific microbiome configuration. Rather than converging within a single microbiome state, including the intermediate Cluster 4, these individuals were distributed across multiple regions of the eubiosis-to-dysbiosis landscape, spanning distinct community configurations (Fig. 5b). This distribution suggests that mild or moderate clinical disease represents a microbiologically heterogeneous phenotype rather than a discrete community type uniformly positioned between periodontal health and severe periodontitis. Thus, comparable levels of mild-to-moderate periodontal destruction may occur within distinct microbial community contexts, once again reflecting that microbiome configuration and clinical disease severity do not have a linear relationship as disease expression is the result of the microbiome intersection with host susceptibility.

Beyond smoking and gingival bleeding, our analysis suggested that systemic factors may also intersect with the subgingival microbiome landscape, although these associations were weaker and less taxon-specific. Individuals with a history of MI were underrepresented in the health-associated state and most common in the terminal dysbiotic state; however, species-level rCCA analyses did not identify direct associations between MI and individual taxa. This pattern may therefore reflect the alignment of MI history with a broader host inflammatory or cardiometabolic phenotype associated with severe periodontitis, rather than a specific MI-associated microbial signature. Consistent with this interpretation, continuous cardiovascular and metabolic measures, including systolic and diastolic blood pressure and fasting blood glucose, showed weaker and more heterogeneous relationships across the landscape (Fig. 3e; Supplementary Fig. S8), suggesting that these systemic factors are not tightly coupled to the eubiosis-to-dysbiosis trajectory that paralleled increasing periodontal severity.

This study leverages a large cross-sectional cohort and an advanced manifold-learning–based computational framework, however several limitations warrant consideration. First, the cross-sectional design restricts causal inference regarding temporal dynamics. Longitudinal sampling will be required to project individual trajectories onto the inferred manifold and track transitions across the principal-tree to validate the temporal dynamics in individual subjects. Second, reliance on 16S rRNA amplicon sequencing limits strain-level and functional resolution. Future deep shotgun metagenomic and transcriptomic studies will be important to determine whether the distinct dysbiotic branches identified here correspond to different functional programs or to alternative taxonomic realizations of similar ecological processes. Third, the selective sampling and site-pooling strategy may obscure fine spatial heterogeneity within the oral cavity. In this respect, however, there is evidence that although there is site to site variability, subgingival microbiome shifts in adults occur at a global, whole mouth, scale [7, 71]. Fourth, the parent case–control sampling frame enriched the cohort for individuals with recent MI and associated cardiometabolic risk factors, potentially limiting the generalizability of the inferred landscape and its host-factor associations. Although the principal-tree architecture was preserved in an analysis restricted to participants without MI, validation in independent cohorts with different demographic and clinical profiles is warranted. Finally, because this cohort was assembled before the widespread use of newer exposures such as electronic cigarettes, the ecological effects of such factors could not be evaluated.

In conclusion, our study revealed that the subgingival microbiome is organized along a continuous but branching ecological landscape spanning eubiosis to dysbiosis. This framework identifies an intermediate destabilized state preceding full dysbiosis and supports the existence of divergent disease-associated trajectories shaped by smoking and gingival inflammation. More broadly, it suggests that periodontitis should be understood not as a single microbial endpoint, but as a set of related ecological states arising from interactions among community structure, host susceptibility, and environmental pressures.

## Methods

### Study cohort

This study leveraged participants from the Buffalo MI–Perio Study, a population-based case– control investigation conducted in Erie and Niagara counties, New York State, between 1997 and 2008 [14, 72, 73]. The parent study was originally designed to examine the association between periodontal disease and incident non-fatal myocardial infarction (MI). A total of 1,355 individuals, aged 35–72 years, were included in the present analysis, comprising 534 subjects discharged alive after an MI event and 821 subjects randomly selected from the underlying community population. Eligible participants met standardized enrollment criteria for the parent study, including the presence of at least six natural teeth to enable comprehensive periodontal assessment. Except for the incident non-fatal myocardial infarction defining case status, individuals with a prior history of coronary heart disease, symptomatic angina, cardiovascular disease requiring dietary or pharmacological management, or cancer were excluded at enrollment. Additional exclusion criteria for the present study included systemic antibiotic use within one month prior to examination. All participants were evaluated at the Center for Preventive Medicine at the University at Buffalo, where calibrated examiners conducted standardized interviews, physical examinations, and full-mouth periodontal assessments. Structured questionnaires were used to collect demographic, behavioral, medical, and oral health-related information, including age, sex, race, education, smoking exposure, alcohol consumption history, dental care and oral hygiene behaviors, physical activity, systemic disease history, and medication use. Physical examinations included anthropometric and blood pressure measurements. Fasting blood specimens were collected by trained phlebotomists for routine biochemical and hematological measurements, including glucose and lipid-related parameters. Detailed descriptions of the original recruitment framework have been reported previously [14, 72]. The study protocol was approved by the Institutional Review Boards of the University at Buffalo and participating hospitals in Erie and Niagara counties (IRB #030-488953). Written informed consent was obtained from all participants prior to enrollment, and all procedures were conducted in accordance with relevant ethical guidelines and regulations governing research involving human subjects.

### Periodontal assessment

Participants completed a comprehensive whole-mouth periodontal examination conducted by calibrated periodontists [14]. The exam used the automated Florida probe and recorded clinical parameters at six sites per tooth for all teeth except third molars. Alveolar crestal height (ACH) was assessed from computer-digitized images of 10 intraoral radiographs. The distance from the cemento-enamel junction to the most coronal aspect of the interproximal alveolar crest was measured at mesial and distal sites of each tooth, excluding canines and third molars due to projection distortion at these positions. Gingival bleeding, an indicator of gingival inflammation also associated with periodontitis, was assessed on the buccal, mesiobuccal, and lingual surfaces of all teeth except third molars. Bleeding was recorded following insertion of a periodontal probe and gentle movement along the gingival sulcus or pocket. Plaque index was assessed by recording visible supragingival plaque as present or absent and was expressed as the percentage of sites with plaque for the whole mouth. Calculus was assessed for each tooth using a three-point scale and summarized as a whole-mouth calculus index, as previously described [74].

### Periodontitis case definitions

Periodontal status was defined using multiple complementary frameworks. Disease was first categorized according to the Centers for Disease Control and Prevention/American Academy of Periodontology (CDC/AAP) surveillance criteria [19]. Mild periodontitis was defined as ≥2 interproximal sites with clinical attachment loss (CAL) ≥3 mm and ≥2 interproximal sites with pocket depth (PD) ≥4 mm, not on the same tooth, or one interproximal site with PD ≥5 mm. Moderate periodontitis was defined as ≥2 interproximal sites with CAL ≥4 mm, not on the same tooth, or ≥2 interproximal sites with PD ≥5 mm, not on the same tooth. Severe periodontitis was defined as ≥2 interproximal sites with CAL ≥6 mm, not on the same tooth, and ≥1 interproximal site with PD ≥5 mm. Participants not meeting any of these criteria were classified as no periodontitis. Because the prevalence of mild periodontitis under this definition was too low in our cohort (3.6%), no periodontitis and mild were combined for subsequent analyses. Periodontal stage was additionally characterized using maximum interproximal CAL in line with the AAP periodontal staging case definitions (2017 World Workshop on the Classification of Periodontal and Peri-Implant Diseases and Conditions [20]). In this stratification, participants were classified as Stage I if they had no interproximal site with CAL ≥3 mm; Stage II if they had at least one interproximal site with CAL of 3–4 mm and no interproximal site with CAL ≥5 mm; and Stage III/IV if they had at least one interproximal site with CAL ≥5 mm. Stage I and Stage II were combined because only 12 participants met the Stage I criterion. In parallel, we applied a custom classification integrating structural destruction and gingival bleeding to distinguish periodontal health, gingivitis, and periodontitis. Periodontitis was defined as the presence of at least one site with both PD ≥5 mm and CAL ≥3 mm. Among participants who did not meet this periodontitis definition, those with gingival bleeding at <20% of sites were classified as periodontally healthy, whereas those with gingival bleeding at ≥20% of sites were classified as having gingivitis. Among individuals with periodontitis, disease severity was further stratified into tertiles based on whole-mouth mean PD to capture gradations in disease burden across the cohort.

### Subgingival plaque sampling and microbiome characterization

Subgingival plaque was sampled from 12 preselected index teeth (#3, 5, 7, 9, 12, 14, 19, 21, 23, 25, 28, and 30) using fine paper points (Johnson & Johnson, East Windsor, NJ, USA), as previously described [75]. If an index tooth was missing, a prespecified substitute tooth was sampled when available. If both the index tooth and its substitute were absent, no sample was collected for that site. Selected teeth were dried with cotton to reduce cross-contamination from saliva or supragingival plaque. One paper point was placed in the gingival pocket or sulcus at the mesiobuccal surface for 10 sec and then transferred to lactated Ringer’s solution. Samples from maxillary and mandibular teeth were pooled and stored separately in liquid nitrogen, and, for the present analysis, pooled into a single vial for each participant prior to DNA extraction. Samples underwent enzymatic lysis consisting of preincubation with achromopeptidase (1000 U/mL) at 37 °C for 10 min, followed by incubation with proteinase K at 56 °C for 30 min. Genomic DNA was subsequently extracted using the QIAamp DNA Mini Kit according to the manufacturer’s instructions. The V3–V4 hypervariable regions of the bacterial 16S rRNA gene were amplified by polymerase chain reaction (PCR) using the universal primer pair 341F (5′-CCTACGGGNGGCWGCAG-3′) and 805R (5′-GACTACHVGGGTATCTAATCC-3′).

Amplicons were purified, indexed, and sequenced on the Illumina MiSeq platform to generate paired-end reads. Raw sequencing data were processed using the DADA2 [76] pipeline implemented in QIIME2 (v2024.10.1) [77]. Quality filtering, denoising, chimera removal, and inference of amplicon sequence variants (ASVs) were conducted following standard parameters. Taxonomic annotation was performed using a two-step approach. ASVs were first aligned against the Human Oral Microbiome Database (HOMD; v15.22) [78] using BLAST [79], and species-level assignments were accepted at ≥99% sequence similarity. Sequences that could not be confidently classified at the species level were further annotated using the Ribosomal Database Project (RDP) classifier [80] to determine the closest reliable taxonomic rank. A total of 68,962,380 sequences (mean ± standard deviation per sample = 50,895 ± 31,513) were retained for downstream analysis. Microbiome composition was characterized using species-level relative abundance subjected to a centered log_2_-ratio (CLR) transformation [81] to account for the compositional structure of the data and reduce the likelihood of spurious correlations [82].

Sequencing data have been deposited in the NCBI Sequence Read Archive under accession number PRJNA1430094.

### Graph-based manifold learning characterization of subgingival microbiome heterogeneity

To characterize population-level heterogeneity in the subgingival microbiome, we implemented an unsupervised machine learning framework to model the latent structure of species-level compositional data. Species-level relative abundance were first transformed using centered log-ratio (CLR) transformation to account for compositional constraints inherent in high-throughput sequencing data [82]. Dimensionality was reduced using the unsupervised iDetect feature selection algorithm [24], which retains species that preserve informative variation and local neighborhood relationships among samples. This step limited the influence of sparse or weakly informative taxa before reconstruction of the low-dimensional microbiome landscape.

Let x*_n_* ɛR^j^ denote the CLR-transformed abundance vector of sample *n* across *j* species. iDetect learns a non-negative feature weight vector wɛR*^j^* by maximizing global variance while preserving local neighborhood structure. This is achieved by optimizing the objective:

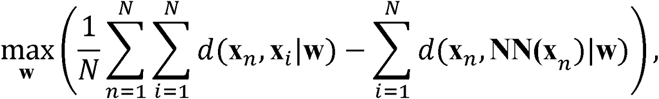

subject to 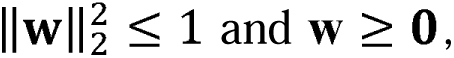 where *d*(,·|**w**) denotes the **w-**weighted block distance, and **NN**(**x***_n_*) represents the nearest neighbor of sample *n*. A probabilistic relaxation of the nearest-neighbor operator is utilized to enable efficient optimization, and an ℓ_1_ penalty (||**w**||_1_≤*λ*) is applied to induce sparsity [24]. The iDetect regularization parameter *λ* was evaluated over a prespecified grid and selected at the elbow of the optimization-objective curve, yielding (λ=15). A total of 599 species with non-zero weights were retained for downstream manifold learning.

Selected species were assembled into a matrix Xɛ R*^D×N^*, which denotes the CLR-transformed abundance matrix comprising *D* selected species across *N* samples. We then applied the DDRTree [25] algorithm to jointly learn a low-dimensional representation and an explicit principal-tree structure. DDRTree estimates an orthogonal projection matrix that maps the high-dimensional abundance matrix **X** into a three-dimensional latent space, yielding embedding coordinates for each sample. This low-dimensional representation was used to reconstruct the subgingival microbiome landscape, in which spatial proximity reflects similarity in community configuration. Within this latent space, a set of representative nodes is introduced and connected through a tree structure estimated using a minimum spanning tree under a reversed graph embedding framework. The algorithm simultaneously optimizes data reconstruction, preservation of global topology, and local assignment of samples to nearby structural nodes. Through this joint optimization, DDRTree learns a principal-tree that traverses the center of the data cloud and approximates the underlying population structure. Importantly, the learned structure adapts to the geometry of the data and can represent gradual transitions or branching configurations when supported by the data. To determine the DDRTree hyperparameters, the local-assignment weight (λ) was fixed at 1, whereas the kernel width (α) and principal-tree regularization parameter (λ) were evaluated over prespecified grids. The final values (α=50 and λ =300) were selected based on the elbow in the trade-off between reconstruction error and principal-tree length.

To identify interpretable microbiome states along the reconstructed microbiome landscape, spectral clustering was applied to the low-dimensional embedding obtained from DDRTree. This graph-based approach is highly effective for partitioning complex, non-convex structures, including branching biological manifolds. To preserve the topology of the learned structure, sample similarity was quantified using geodesic distance along the inferred principal-tree, defined as the shortest path length between sample projections on the tree, rather than Euclidean distance in the embedding space. The optimal number of clusters was determined based on gap statistics and average silhouette width. Cluster assignments were subsequently used to define discrete microbiome states and evaluate their associations with host clinical and demographic variables.

### Regularized canonical correlation analysis

To investigate the association between microbiome composition and host metadata, we performed regularized canonical correlation analysis (rCCA) using the mixOmics [83] package in R. rCCA is an unsupervised multivariate technique that uncovers relationships between two sets of variables by projecting them into a shared latent space defined by canonical variates. Two complementary analyses were conducted using the same rCCA framework and parameter optimization strategy. First, microbiome clusters derived from the inferred microbiome landscape were one-hot encoded and analyzed against host metadata to evaluate associations between community-level cluster structure and host factors. Second, species-level abundance data were analyzed against the same metadata to characterize taxon-specific associations. For this analysis, microbiome data were filtered to exclude species present in fewer than 10% of samples (>90% zero counts) to reduce sparsity and enhance signal robustness. Remaining species were normalized using a robust centered log-ratio transformation to account for compositional constraints. Host and environmental metadata were preprocessed using the Yeo– Johnson transformation to stabilize variance and approximate normality. For both analyses, regularization parameters (λ□, λ□) were optimized using 5-fold cross-validation to balance model complexity and generalizability and to mitigate multicollinearity in high-dimensional datasets. The number of canonical variates retained was determined independently for each model based on the elbow point of the cumulative shared variance curve. Accordingly, three canonical variates were retained for the cluster-level analysis, whereas eight canonical variates were retained for the species-level analysis, because the species-level analysis involved greater data dimensionality and complexity. Canonical loadings derived from the retained variates were used to interpret associations between microbiome features and host variables. Associations between microbiome features and host variables were quantified as the inner product of their corresponding canonical loading vectors within the retained canonical subspace.

### Statistical analyses

Differences in host demographic and clinical characteristics between groups were assessed using one-way analysis of variance (ANOVA) for continuous variables and χ² tests for categorical variables. Candidate covariates for the adjusted diversity analyses were identified from demographic, lifestyle, and cardiometabolic variables that differed across periodontal groups under at least one periodontal disease definition (*P□*<□0.05). Pairwise Spearman’s correlation coefficients were calculated among these variables, and *P* values were adjusted using the Bonferroni correction for multiple testing (Supplementary Fig. S12). The final covariate set was selected based on the observed correlation structure together with biological relevance and the design of the parent study. The resulting adjustment set for the subsequent alpha-and beta-diversity analyses comprised age, sex, years of education, history of myocardial infarction, current smoking status, lifetime smoking exposure in pack-years, lifetime alcohol consumption, annual dental visits, and daily flossing. Species-level alpha diversity (within-sample richness) was quantified using the Chao1 index [84] following rarefaction to 1,951 reads per sample, corresponding to the minimum sequencing depth. Rarefaction was repeated 1,000 times, and the average value was reported. Differences in alpha diversity among periodontal groups were evaluated using analysis of covariance (ANCOVA). Beta diversity (between-sample differences in community composition) was quantified using Euclidean distances between species-level centered log-ratio (CLR)-transformed abundance profiles (i.e., Aitchison distances) and visualized using principal component analysis (PCA). Differences in community composition among periodontal groups were evaluated using permutational multivariate analysis of variance (PERMANOVA). Covariate adjustment was applied to both the ANCOVA and PERMANOVA models. To quantify sample-level dysbiosis, the subgingival microbiome dysbiosis index (SMDI) was calculated as described previously, as the difference between the mean CLR abundance of periodontitis-associated species and that of health-associated species [27]. Of the 49 species in the published signature, 47 were represented in our dataset and included in the calculation; *Peptostreptococcaceae [XI][G-4] sp. oral taxon 103* and *Streptococcus infantis* were not represented. Differentially abundant taxa across microbiome clusters were identified using linear discriminant analysis effect size (LEfSe) to detect taxa enriched in specific clusters or cluster groupings. Taxa were annotated as health-, gingivitis-, or periodontitis-associated primarily according to a previous species-level synthesis by Diaz et al. [26], which integrated evidence from high-throughput sequencing studies of subgingival microbiome shifts across periodontal conditions. For taxa not categorized in that synthesis, annotations were supplemented using the updated meta-analysis and unified reanalysis by Abusleme et al. [6], which defined microbial signatures of health, gingivitis, and periodontitis using harmonized sequence processing and taxonomic classification across publicly available datasets. Using the results of Abusleme et al., taxa were assigned to a periodontal condition based on LEfSe-defined enrichment with an linear discriminant analysis (LDA) score threshold >3. For analyses involving multiple comparisons, *P* values were adjusted within each analysis using the Bonferroni method, and adjusted *P* < 0.05 was considered statistically significant. All statistical analyses were conducted in MATLAB (R2025b) unless otherwise indicated.

## List of abbreviations

AAP: American Academy of Periodontology
ACH: Alveolar crestal height
ANCOVA: Analysis of covariance
ANOVA: Analysis of variance
ASV: Amplicon sequence variant
CAL: Clinical attachment loss
CDC: Centers for Disease Control and Prevention
CLR: Centered log_2_-ratio
DDRTree: Discriminative dimensionality reduction tree
HOMD: Human Oral Microbiome Database
LEfSe: Linear discriminant analysis effect size
MI: Myocardial infarction
PCA: Principal component analysis
PCR: Polymerase chain reaction
PD: Pocket depth
PERMANOVA: Permutational multivariate analysis of variance
rCCA: Regularized canonical correlation analysis
RDP: Ribosomal Database Project
SMDI: Subgingival microbial dysbiosis index

## Declarations

### Ethics approval and consent to participate

The study protocol was approved by the Institutional Review Boards of the University at Buffalo and participating hospitals in Erie and Niagara counties (IRB #030-488953). Written informed consent to participate was obtained from all participants.

### Consent for publication

Not applicable.

### Data availability

The microbiome sequence data have been deposited in the NCBI Sequence Read Archive under BioProject accession PRJNA1430094. Associated demographic and clinical metadata are available in the linked BioSample records. The custom MATLAB code for manifold reconstruction, clustering, and trajectory analysis, together with the R code for rCCA, is publicly available at https://github.com/liluacrobat/Subgingival-Ecological-Landscape.

### Authors’ contributions

L.L. and P.I.D. led the conception and design of the study, with contributions from all co-authors. M.H. and H.H. contributed substantially to the development of the analytical approach. J.W.W., K.L.F., R.J.G., Y.S. and M.J.B. contributed to data acquisition and methodology development. L.L. performed the analyses, with M.H. conducting the regularized canonical correlation analysis (rCCA). L.L., M.H., H.H., and P.I.D. interpreted the results. L.L. and P.I.D. wrote the original manuscript. All authors contributed to manuscript review and editing. All surviving authors confirm that R.J.G. met the authorship criteria and agree to his inclusion as an author.

## References

1. Löe, H., et al., The natural history of periodontal disease in man. Study design and baseline data. J Periodontal Res, 1978. 13(6): p. 550–62.

2. Löe, H., et al., Natural history of periodontal disease in man. Rapid, moderate and no loss of attachment in Sri Lankan laborers 14 to 46 years of age. J Clin Periodontol, 1986. 13(5): p. 431–45.

3. Genco, R.J. and W.S. Borgnakke, Risk factors for periodontal disease. Periodontol 2000, 2013. 62(1): p. 59–94.

4. Eke, P.I., W.S. Borgnakke, and R.J. Genco, Recent epidemiologic trends in periodontitis in the USA. Periodontol 2000, 2020. 82(1): p. 257–267.

5. Socransky, S.S., et al., Microbial complexes in subgingival plaque. J Clin Periodontol, 1998. 25(2): p. 134–44.

6. Abusleme, L., et al., Microbial signatures of health, gingivitis, and periodontitis. Periodontol 2000, 2021. 86(1): p. 57–78.

7. Griffen, A.L., et al., Distinct and complex bacterial profiles in human periodontitis and health revealed by 16S pyrosequencing. ISME J, 2012. 6(6): p. 1176–85.

8. Camelo-Castillo, A.J., et al., Subgingival microbiota in health compared to periodontitis and the influence of smoking. Front Microbiol, 2015. 6: p. 119.

9. Ganesan, S.M., et al., A tale of two risks: smoking, diabetes and the subgingival microbiome. ISME J, 2017. 11(9): p. 2075–2089.

10. Qu, H. and S. Zhang, Association of cardiovascular health and periodontitis: a population-based study. BMC Public Health, 2024. 24(1): p. 438.

11. Li, Y., et al., Metabolic syndrome exacerbates inflammation and bone loss in periodontitis. J Dent Res, 2015. 94(2): p. 362–70.

12. Wang, J., et al., Alcohol consumption and risk of periodontitis: a meta-analysis. J Clin Periodontol, 2016. 43(7): p. 572–83.

13. Marruganti, C., et al., Adherence to Mediterranean diet, physical activity level, and severity of periodontitis: results from a university-based cross-sectional study. J Periodontol, 2022. 93(8): p. 1218–1232.

14. Andriankaja, O.M., et al., The use of different measurements and definitions of periodontal disease in the study of the association between periodontal disease and risk of myocardial infarction. J Periodontol, 2006. 77(6): p. 1067–73.

15. Amaral Cda, S., et al., Evaluation of the subgingival microbiota of alcoholic and non-alcoholic individuals. J Dent, 2011. 39(11): p. 729–38.

16. Khocht, A., et al., Metabolomic profiles of obesity and subgingival microbiome in periodontally healthy individuals: a cross-sectional study. J Clin Periodontol, 2023. 50(11): p. 1455–1466.

17. Shah, S.A., et al., The making of a miscreant: tobacco smoke and the creation of pathogen-rich biofilms. NPJ Biofilms Microbiomes, 2017. 3: p. 26.

18. Li, L., et al., Computational approach to modeling microbiome landscapes associated with chronic human disease progression. PLoS Comput Biol, 2022. 18(8): p. e1010373.

19. Eke, P.I., et al., Update of the case definitions for population-based surveillance of periodontitis. J Periodontol, 2012. 83(12): p. 1449–54.

20. Tonetti, M.S., H. Greenwell, and K.S. Kornman, Staging and grading of periodontitis: framework and proposal of a new classification and case definition. J Clin Periodontol, 2018. 45 Suppl 20: p. S149–s161.

21. Genco, R.J., et al., The subgingival microbiome relationship to periodontal disease in older women. J Dent Res, 2019. 98(9): p. 975–984.

22. Iniesta, M., et al., Subgingival microbiome in periodontal health, gingivitis and different stages of periodontitis. J Clin Periodontol, 2023. 50(7): p. 905–920.

23. Abusleme, L., et al., The subgingival microbiome in health and periodontitis and its relationship with community biomass and inflammation. ISME J, 2013. 7(5): p. 1016–25.

24. Yao, J., et al., Feature selection for unsupervised learning through local learning. Pattern Recognition Letters, 2015. 53: p. 100–107.

25. Mao, Q., et al. Dimensionality reduction via graph structure learning. in Proceedings of the 21th ACM SIGKDD International Conference on Knowledge Discovery and Data Mining. 2015.

26. Diaz, P.I., A. Hoare, and B.Y. Hong, Subgingival microbiome shifts and community dynamics in periodontal diseases. J Calif Dent Assoc, 2016. 44(7): p. 421–35.

27. Chen, T., P.D. Marsh, and N.N. Al-Hebshi, SMDI: an index for measuring subgingival microbial dysbiosis. J Dent Res, 2022. 101(3): p. 331–338.

28. Gomes-Filho, I.S., et al., Severe and moderate periodontitis are associated with acute myocardial infarction. J Periodontol, 2020. 91(11): p. 1444–1452.

29. Thapa, S. and F. Wei, Association between high serum total cholesterol and periodontitis: national health and nutrition examination survey 2011 to 2012 study of American adults. J Periodontol, 2016. 87(11): p. 1286–1294.

30. Haber, J., et al., Evidence for cigarette smoking as a major risk factor for periodontitis. J Periodontol, 1993. 64(1): p. 16–23.

31. Baumeister, S.E., et al., Alcohol consumption, risk of periodontitis and change of periodontal parameters in a population-based cohort study. J Clin Periodontol, 2025. 52(7): p. 1024–1031.

32. Greene, J.C., Oral hygiene and periodontal disease. American Journal of Public Health and the Nations Health, 1963. 53(6): p. 913–922.

33. Zadik, Y., et al., Periodontal disease might be associated even with impaired fasting glucose. Br Dent J, 2010. 208(10): p. E20.

34. Preshaw, P.M. and S.M. Bissett, Periodontitis and diabetes. British dental journal, 2019. 227(7): p. 577–584.

35. Suvan, J.E., et al., Association between overweight/obesity and increased risk of periodontitis. J Clin Periodontol, 2015. 42(8): p. 733–739.

36. Haffajee, A.D. and S.S. Socransky, Relationship of cigarette smoking to the subgingival microbiota. J Clin Periodontol, 2001. 28(5): p. 377–88.

37. Mason, M.R., et al., The subgingival microbiome of clinically healthy current and never smokers. ISME J, 2015. 9(1): p. 268–72.

38. Li, L., et al., Effect of an intensive antiplaque regimen on microbiome outcomes after nonsurgical periodontal therapy. J Periodontol, 2025. 96(3): p. 241–254.

39. Hagenfeld, D., et al., Long-term changes in the subgingival microbiota in patients with stage III-IV periodontitis treated by mechanical therapy and adjunctive systemic antibiotics: a secondary analysis of a randomized controlled trial. J Clin Periodontol, 2023. 50(8): p. 1101–1112.

40. Kumar, P.S., et al., Subgingival host-microbial interactions in hyperglycemic individuals. J Dent Res, 2020. 99(6): p. 650–657.

41. Bizzarro, S., et al., Microbial profiles at baseline and not the use of antibiotics determine the clinical outcome of the treatment of chronic periodontitis. Sci Rep, 2016. 6: p. 20205.

42. Duran-Pinedo, A.E., et al., Subgingival host-microbiome metatranscriptomic changes following scaling and root planing in grade II/III periodontitis. J Clin Periodontol, 2023. 50(3): p. 316–330.

43. Chen, C., et al., Oral microbiota of periodontal health and disease and their changes after nonsurgical periodontal therapy. ISME J, 2018. 12(5): p. 1210–1224.

44. Hajishengallis, G., The inflammophilic character of the periodontitis-associated microbiota. Mol Oral Microbiol, 2014. 29(6): p. 248–57.

45. Hajishengallis, G. and R.J. Lamont, Beyond the red complex and into more complexity: the polymicrobial synergy and dysbiosis (PSD) model of periodontal disease etiology. Mol Oral Microbiol, 2012. 27(6): p. 409–19.

46. Qin, X., Y. Zhao, and Y. Guo, Periodontal disease and myocardial infarction risk: a meta-analysis of cohort studies. Am J Emerg Med, 2021. 48: p. 103–109.

47. Sanz, M., et al., Periodontitis and cardiovascular diseases. Consensus report. Glob Heart, 2020. 15(1): p. 1.

48. Rutger Persson, G., et al., Chronic periodontitis, a significant relationship with acute myocardial infarction. Eur Heart J, 2003. 24(23): p. 2108–15.

49. Kolenbrander, P.E., et al., Oral multispecies biofilm development and the key role of cell-cell distance. Nat Rev Microbiol, 2010. 8(7): p. 471–80.

50. Zijnge, V., et al., Oral biofilm architecture on natural teeth. PLoS One, 2010. 5(2): p. e9321.

51. Zhang, M., M. Whiteley, and G.R. Lewin, Polymicrobial interactions of oral microbiota: a historical review and current perspective. mBio, 2022. 13(3): p. e0023522.

52. Diaz, P.I., P.S. Zilm, and A.H. Rogers, *Fusobacterium nucleatum* supports the growth of *Porphyromonas gingivalis* in oxygenated and carbon-dioxide-depleted environments. Microbiology (Reading), 2002. 148(Pt 2): p. 467–472.

53. Sakanaka, A., et al., *Fusobacterium nucleatum* metabolically integrates commensals and pathogens in oral biofilms. mSystems, 2022. 7(4): p. e0017022.

54. Albaghdadi, S.Z., et al., In vitro characterization of biofilm formation in *Prevotella* species. Front Oral Health, 2021. 2: p. 724194.

55. Kolenbrander, P.E., R.N. Andersen, and L.V. Holdeman, Coaggregation of oral *Bacteroides* species with other bacteria: central role in coaggregation bridges and competitions. Infect Immun, 1985. 48(3): p. 741–6.

56. Guan, S.M., et al., Degradation of human hemoglobin by *Prevotella intermedia*. Anaerobe, 2006. 12(5-6): p. 279–82.

57. Takahashi, N. and T. Yamada, Pathways for amino acid metabolism by *Prevotella intermedia* and *Prevotella nigrescens*. Oral Microbiol Immunol, 2000. 15(2): p. 96–102.

58. Mark Welch, J.L., et al., Biogeography of a human oral microbiome at the micron scale. Proc Natl Acad Sci U S A, 2016. 113(6): p. E791–800.

59. Bergström, J. and L. Boström, Tobacco smoking and periodontal hemorrhagic responsiveness. J Clin Periodontol, 2001. 28(7): p. 680–5.

60. Morozumi, T., et al., Smoking cessation increases gingival blood flow and gingival crevicular fluid. J Clin Periodontol, 2004. 31(4): p. 267–72.

61. Dietrich, T., J.P. Bernimoulin, and R.J. Glynn, The effect of cigarette smoking on gingival bleeding. J Periodontol, 2004. 75(1): p. 16–22.

62. Bagaitkar, J., et al., Tobacco-induced alterations to *Porphyromonas gingivalis*-host interactions. Environ Microbiol, 2009. 11(5): p. 1242–53.

63. Olsen, I. and G. Hajishengallis, Major neutrophil functions subverted by *Porphyromonas gingivalis*. J Oral Microbiol, 2016. 8: p. 30936.

64. Zenobia, C. and G. Hajishengallis, *Porphyromonas gingivalis* virulence factors involved in subversion of leukocytes and microbial dysbiosis. Virulence, 2015. 6(3): p. 236–43.

65. Hajishengallis, G., R.P. Darveau, and M.A. Curtis, The keystone-pathogen hypothesis. Nat Rev Microbiol, 2012. 10(10): p. 717–25.

66. Nibali, L., et al., What is the heritability of periodontitis? A systematic review. J Dent Res, 2019. 98(6): p. 632–641.

67. Schaefer, A.S., et al., Genetic risk variants implicate impaired maintenance and repair of periodontal tissues as causal for periodontitis-a synthesis of recent findings. Periodontol 2000, 2025.

68. Dayan, S., et al., Oral epithelial overexpression of IL-1alpha causes periodontal disease. J Dent Res, 2004. 83(10): p. 786–90.

69. Maekawa, T., et al., *Porphyromonas gingivalis* manipulates complement and TLR signaling to uncouple bacterial clearance from inflammation and promote dysbiosis. Cell Host Microbe, 2014. 15(6): p. 768–78.

70. Dutzan, N., et al., A dysbiotic microbiome triggers T(H)17 cells to mediate oral mucosal immunopathology in mice and humans. Sci Transl Med, 2018. 10(463).

71. Dabdoub, S.M., S.M. Ganesan, and P.S. Kumar, Comparative metagenomics reveals taxonomically idiosyncratic yet functionally congruent communities in periodontitis. Sci Rep, 2016. 6: p. 38993.

72. Andriankaja, O.M., et al., Periodontal disease and risk of myocardial infarction: the role of gender and smoking. Eur J Epidemiol, 2007. 22(10): p. 699–705.

73. Andriankaja, O., et al., Association between periodontal pathogens and risk of nonfatal myocardial infarction. Community Dent Oral Epidemiol, 2011. 39(2): p. 177–85.

74. Machtei, E.E., et al., The rate of periodontal attachment loss in subjects with established periodontitis. J Periodontol, 1993. 64(8): p. 713–8.

75. Brennan-Calanan, R.M., et al., Osteoporosis and oral infection: independent risk factors for oral bone loss. J Dent Res, 2008. 87(4): p. 323–7.

76. Callahan, B.J., et al., DADA2: high-resolution sample inference from Illumina amplicon data. Nat Methods, 2016. 13(7): p. 581–3.

77. Bolyen, E., et al., Reproducible, interactive, scalable and extensible microbiome data science using QIIME 2. Nat Biotechnol, 2019. 37(8): p. 852–857.

78. Dewhirst, F.E., et al., The human oral microbiome. J Bacteriol, 2010. 192(19): p. 5002–17.

79. Camacho, C., et al., BLAST+: architecture and applications. BMC Bioinformatics, 2009. 10: p. 421.

80. Maidak, B.L., et al., The ribosomal database project. Nucleic Acids Res, 1994. 22(17): p. 3485–7.

81. Greenacre, M., Compositional data analysis in practice. 2018: Chapman and Hall/CRC.

82. Gloor, G.B., et al., Microbiome datasets are compositional: and this is not optional. Front Microbiol, 2017. 8: p. 2224.

83. Rohart, F., et al., mixOmics: an R package for ‘omics feature selection and multiple data integration. PLoS Comput Biol, 2017. 13(11): p. e1005752.

84. Chao, A. and C.-H. Chiu, Species richness: estimation and comparison. Wiley StatsRef: statistics reference online, 2016. 1(26): p. 10.1002.

