## Supplementary Figures for "A population-scale landscape of the subgingival microbiome reveals divergent routes to periodontal dysbiosis"

**Supplementary Figure S1. Species-level alpha and beta diversity of the subgingival microbiome in relation to AAP periodontal stage case definitions.** (a) Species-level alpha diversity of AAP stage groups. Stages I/II included subjects with interdental clinical attachment loss (CAL) ≤ 4 mm, whereas Stages III/IV included subjects with interdental CAL ≥ 5 mm. (b) Beta diversity via principal components analysis (PCA) depicting Euclidean distances among AAP staging groups. Statistical significance for alpha diversity was assessed using ANCOVA and for beta diversity using PERMANOVA, respectively, with adjustment for age, sex, years of education, history of myocardial infarction, current smoking status, lifetime smoking exposure in pack-years, lifetime alcohol consumption, annual dental visits, and daily flossing. ** = *P* < 0.01.

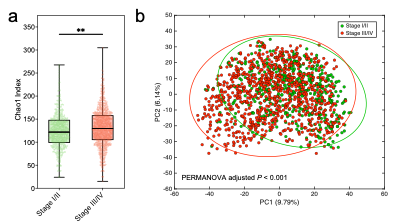

**Supplementary Figure S2. Beta diversity differences between periodontal healthy subjects and those with severe periodontitis.** (a) Principal components analysis (PCA) of beta diversity based on Euclidean distances comparing none/mild versus severe groups defined by CDC/AAP criteria. (b) PCA of beta diversity based on Euclidean distances comparing health and periodontitis with high mean pocket depth (PD) using the custom periodontal classification. Statistical significance was assessed using conditioned PERMANOVA. Models were adjusted for history of myocardial infarction, current smoker, annual dental visits, and definition-specific covariates: age and years of education for the CDC/AAP grouping, and daily flossing for the custom periodontal classification.

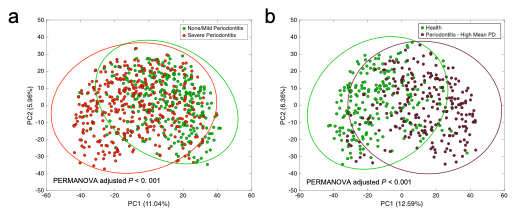

**Supplementary Figure S3. Sensitivity analysis of the inferred microbiome structure according to MI history.** (a) Tree-like trajectory inferred from the full dataset, with samples color-coded by MI status. (b) Tree-like trajectory reconstructed using only samples from control subjects.

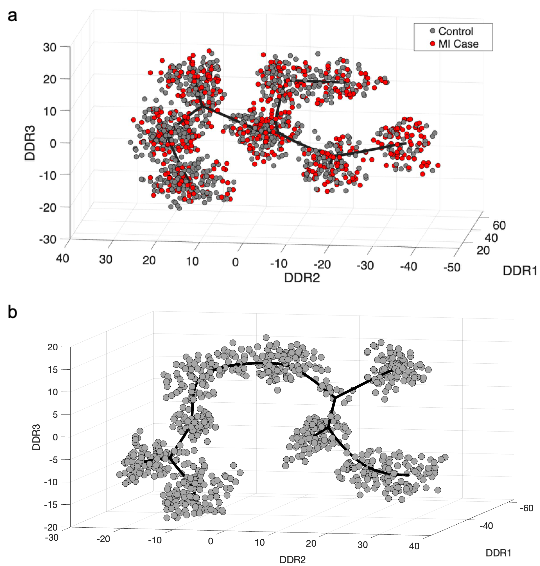

**Supplementary Figure S4. Heatmap showing the relative abundance of cluster-specific core species.** Core species are defined as those with an average relative abundance >1% and a prevalence > 70% within each cluster.

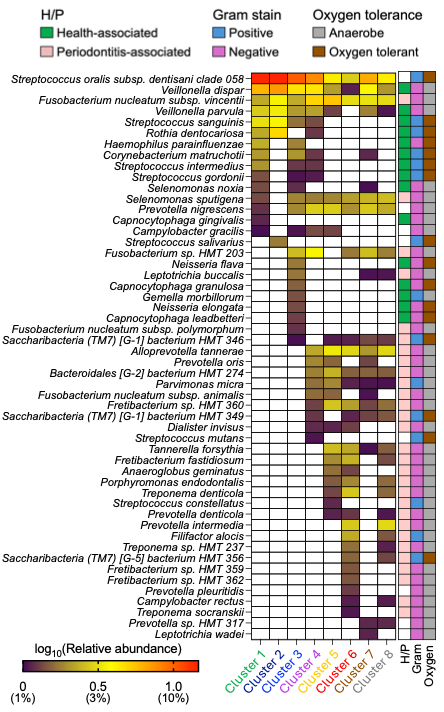

**Supplementary Figure S5. Distributions of periodontal clinical parameters across microbiome states.** Box plots show periodontal clinical measures that differed significantly across microbiome clusters. Statistical significance was assessed using one-way ANOVA with Bonferroni correction for pairwise comparisons. Letters above the boxes indicate pairwise comparison results; clusters sharing at least one letter are not significantly different, whereas clusters with no shared letters differ significantly (*P* < 0.05).

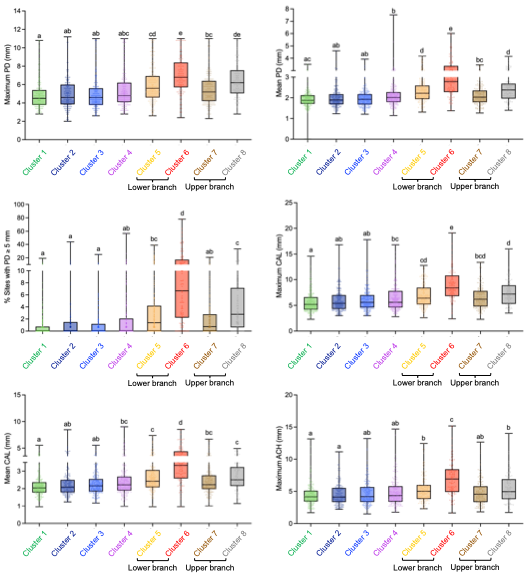

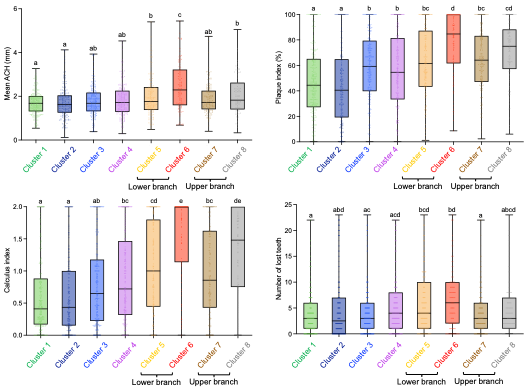

**Supplementary Figure S6. Distributions of host-related variables other than periodontal clinical parameters across microbiome states.** Box plots show continuous variables, and bar plots show the prevalence of categorical variables across microbiome clusters. For continuous variables, statistical significance was first assessed using one-way ANOVA, followed by post hoc pairwise comparisons. For categorical variables, pairwise differences between clusters were assessed using chi-squared tests. Bonferroni correction was applied to account for multiple comparisons. Letters above the plots indicate pairwise comparison results; clusters sharing at least one letter are not significantly different, whereas clusters with no shared letters differ significantly (*P* < 0.05).

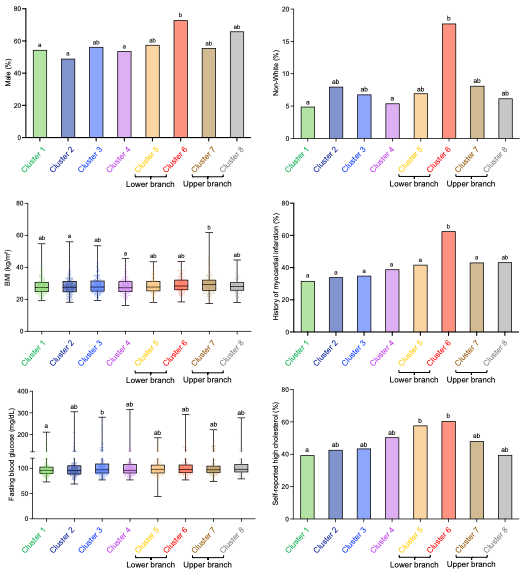

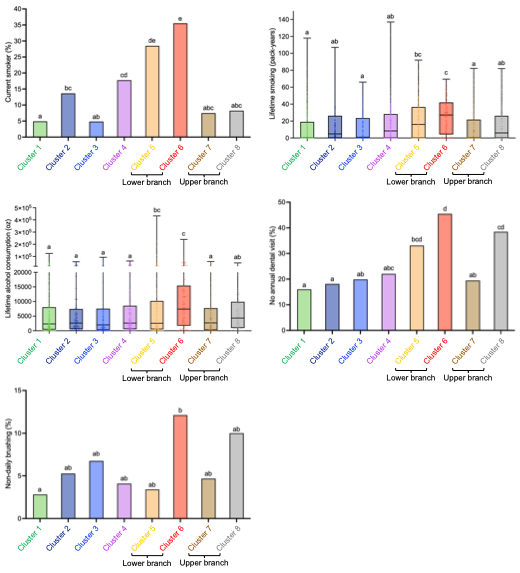

**Supplementary Figure S7. Associations between microbial taxa and host factors across microbiome states.** Heatmap showing association scores between individual taxa and host factors based on regularized canonical correlation analysis (rCCA). Only associations with an absolute value ≥ 0.15 are displayed.

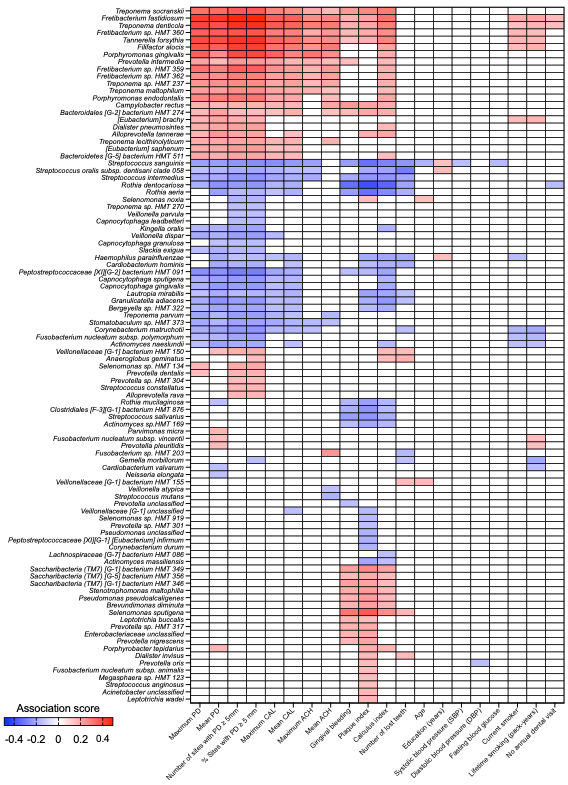

**Supplementary Figure S8. Species peaking in Cluster 4 representing an intermediate ecological configuration between eubiotic and branch-specific dysbiotic states.** Box plots show the relative abundance of species that reached their highest abundance in Cluster 4 across four ecological states: eubiotic states (Clusters 1–3), Cluster 4, the lower branch (Clusters 5 and 6), and the upper branch (Clusters 7 and 8). Statistical significance was assessed using the Kruskal–Wallis test followed by Bonferroni-corrected post hoc pairwise comparisons. * = *P* < 0.05, ** = *P* < 0.01, *** = *P* < 0.001.

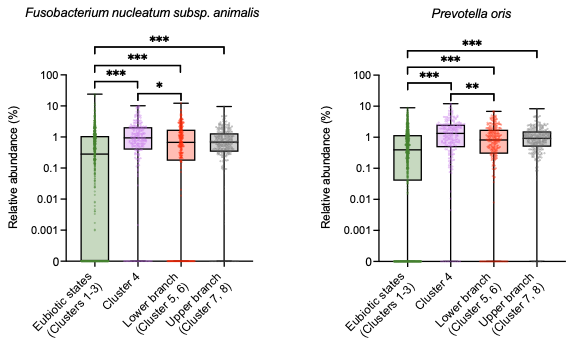

**Supplementary Fig. S9. Moderate Pearson correlations between subgingival microbial dysbiosis index and clinical periodontal measures reveal discordant microbiome–clinical phenotypes.** Scatter plots show Pearson correlations between the subgingival microbial dysbiosis index (SMDI) and mean PD or maximum CAL. Green points indicate individuals with high dysbiosis but low clinical damage, defined as SMDI above the 85th percentile and the corresponding clinical measure below the 15th percentile. Red points indicate individuals with low dysbiosis but high clinical damage, defined as SMDI below the 15th percentile and the corresponding clinical measure above the 85th percentile.

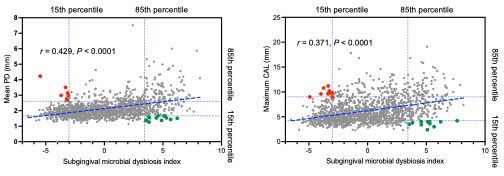

**Supplementary Figure S10. Microbial profiles of health and periodontitis within Cluster 1.** Pie plots show the relative abundance of the top 20 most abundant genera among individuals with periodontitis and high mean probing depth (Periodontitis–High Mean PD) within Cluster 1. Each pie plot represents one individual.

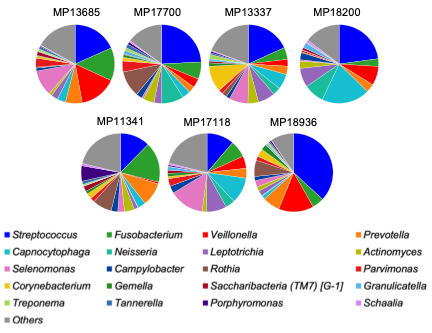

**Supplementary Figure S11. Microbial profiles of clinically healthy individuals and Cluster 6-enriched species across periodontal phenotypes within Cluster 6.** (a) Pie plots show the relative abundance of the 20 most abundant genera among clinically healthy individuals within Cluster 6. Each pie plot represents one individual. (b) Relative abundance of Cluster 6-enriched pathobionts across Cluster 6 subjects with distant periodontal phenotypes. These species were identified by LEfSe as the taxa most strongly associated with Cluster 6 when compared to the remaining clusters. No significant differences were detected across phenotype groups (*P* > 0.05; Kruskal–Wallis test). ns, not significant.

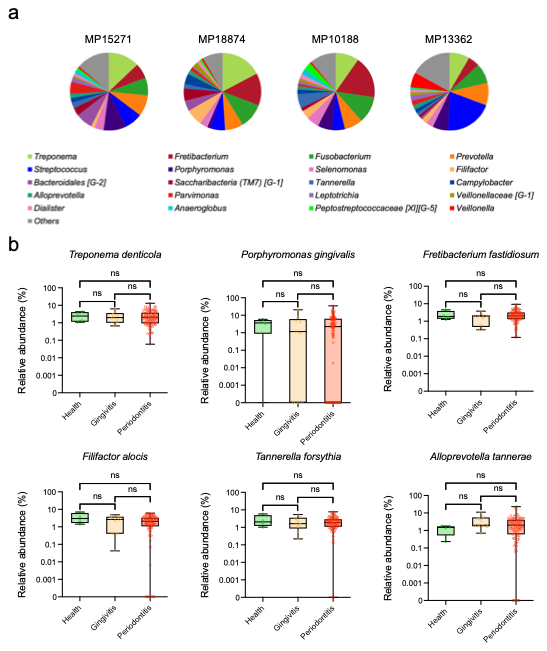

**Supplementary Figure S12. Correlation structure among candidate covariates.** Heatmap showing pairwise Spearman’s correlations among demographic, lifestyle, and cardiometabolic variables that differed across periodontal groups under at least one periodontal disease definition. Bonferroni correction was applied to account for multiple comparisons. Correlations with Bonferroni-adjusted *P* ≥ 0.05 were set to zero and are shown in white. Red and blue indicate positive and negative correlations, respectively, with color intensity representing correlation magnitude. Variables were ordered to group related domains.

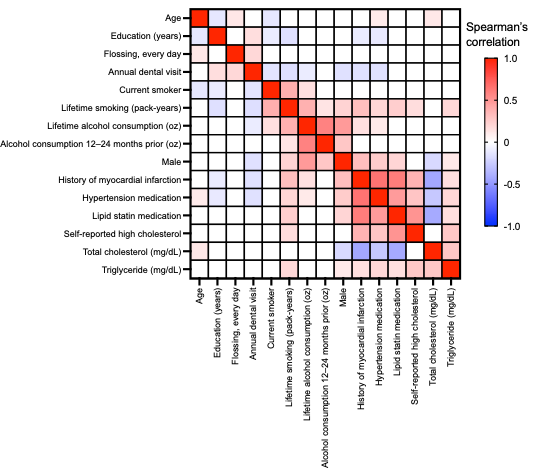
