## Supplementary Table S1 for "A population-scale landscape of the subgingival microbiome reveals divergent routes to periodontal dysbiosis"

**Table 1.** Characteristics of participants by periodontal health status according to the CDC/AAP case definition.

|  | **All**  **(n=1355)** | **Periodontal health (CDC/AAP Categories)** | | | | | |
| --- | --- | --- | --- | --- | --- | --- | --- |
|  |  | **None/Mild**  **(n=311)** | | **Moderate**  **(n=675)** | | **Severe**  **(n=369)** | **Adjusted P-value** |
| **Demographics** |  | | | | | | |
| Age (years), mean (SD) | 55.74 (9.51) | | 53.57 (9.35) | | 56.27 (9.82) | 56.59 (8.78) | <0.01 |
| Male (%) | 56.61 | | 37.94 | | 56.3 | 72.9 | <0.01 |
| Education (years), mean (SD) | 13.70 (2.32) | | 14.11 (2.31) | | 13.73 (2.32) | 13.29 (2.27) | <0.01 |
| Non-White (%) | 7.45 | | 5.47 | | 6.81 | 10.3 | 1.00 |
| BMI (kg/m2), mean (SD) | 28.63 (5.40) | | 28.00 (5.13) | | 28.65 (5.32) | 29.13 (5.71) | 0.98 |
| **Lifestyle and behavioral factors** |  | | | | | | |
| Current smoker (%) | 13.82 | | 7.1 | | 12.31 | 22.22 | <0.01 |
| Lifetime smoking (pack-years), mean (SD) | 15.60 (20.32) | | 7.31 (15.37) | | 14.80 (19.94) | 24.04 (21.46) | <0.01 |
| Lifetime alcohol consumption (oz),  mean (SD) | 8340.23 (20406.58) | | 4992.13 (14129.36) | | 7846.16 (20267.69) | 12090.63 (24266.23) | <0.01 |
| Alcohol consumption during the 12-month  period 12–24 months prior (oz), mean (SD) | 135.42 (315.28) | | 95.90 (301.11) | | 125.60 (259.02) | 186.36 (403.06) | 0.02 |
| Annual dental visit, (%) | 75.94 | | 84.56 | | 77.52 | 65.61 | <0.01 |
| Tooth brushing, every day (%) | 94.52 | | 95.29 | | 95.45 | 92.13 | 1.00 |
| Flossing, every day (%) | 22.12 | | 18.67 | | 23.26 | 23.05 | 1.00 |
| Exercise hours per week, mean (SD) | 5.12 (1.78) | | 5.13 (1.70) | | 5.18 (1.73) | 5.00 (1.92) | 1.00 |
| **Medical history and medication use** |  | | | | | | |
| History of myocardial infarction (%) | 39.41 | | 22.83 | | 37.19 | 57.45 | <0.01 |
| Self-reported hypertension (%) | 36.1 | | 28.8 | | 36.94 | 40.71 | 0.18 |
| Self-reported diabetes (%) | 9.81 | | 6.8 | | 9.97 | 12.09 | 1.00 |
| Systolic blood pressure (SBP) (mmHg),  mean (SD) | 118.27 (14.91) | | 117.66 (14.99) | | 117.90 (14.65) | 119.44 (15.31) | 1.00 |
| Diastolic blood pressure (DBP) (mmHg),  mean (SD) | 72.26 (9.25) | | 72.26 (9.49) | | 71.63 (8.90) | 73.41 (9.58) | 0.47 |
| Hypertension medication (%) | 47.93 | | 34.73 | | 48.74 | 57.61 | <0.01 |
| Lipid statin medication (%) | 28.71 | | 18.01 | | 26.96 | 40.92 | <0.01 |
| Diabetes medication (%) | 7.68 | | 4.82 | | 8.15 | 9.21 | 1.00 |
| Steroid medication (%) | 1.62 | | 1.61 | | 1.48 | 1.9 | 1.00 |
| **Blood glucose and lipid measures** |  | | | | | | |
| Fasting blood glucose (mg/dL), mean (SD) | 104.07 (29.08) | | 100.32 (23.99) | | 104.73 (31.47) | 106.04 (28.14) | 1.00 |
| Self-reported high cholesterol (%) | 47.01 | | 34.84 | | 47.31 | 56.98 | <0.01 |
| Total cholesterol (mg/dL), mean (SD) | 201.49 (43.09) | | 205.56 (39.00) | | 202.98 (43.37) | 195.24 (45.27) | 0.17 |
| Triglyceride (mg/dL), mean (SD) | 146.83 (90.58) | | 130.09 (79.06) | | 151.47 (93.21) | 154.32 (94.17) | 0.04 |
| **Periodontal clinical parameters** |  | | | | | | |
| Number of lost teeth, mean (SD) | 4.97 (5.48) | | 4.02 (5.50) | | 5.15 (5.65) | 5.43 (5.07) | 0.07 |
| Pocket depth (PD), mean (SD) |  | | | | | | |
| Whole mouth mean (mm) | 2.15 (0.60) | | 1.78 (0.42) | | 2.02 (0.34) | 2.69 (0.72) | <0.01 |
| Maximum (Deepest site) (mm) | 5.48 (1.89) | | 3.92 (0.98) | | 5.09 (1.35) | 7.50 (1.61) | <0.01 |
| Clinical attachment loss (CAL), mean (SD) |  | | | | | | |
| Whole mouth mean (mm) | 2.51 (1.01) | | 1.75 (0.30) | | 2.33 (0.59) | 3.46 (1.25) | <0.01 |
| Maximum (mm) | 6.51 (2.46) | | 4.22 (0.93) | | 6.06 (1.58) | 9.25 (2.14) | <0.01 |
| Alveolar crestal height (ACH), mean (SD) |  | | | | | | |
| Whole mouth mean (mm) | 1.87 (0.78) | | 1.43 (0.47) | | 1.71 (0.56) | 2.49 (0.93) | <0.01 |
| Maximum (mm) | 5.00 (2.09) | | 3.56 (1.01) | | 4.69 (1.58) | 6.71 (2.38) | <0.01 |
| Number of sites with PD ≥ 5 mm, mean (SD) | 3.68 (8.37) | | 0.08 (0.32) | | 1.21 (1.98) | 11.24 (13.08) | <0.01 |
| % Sites with PD ≥ 5 mm, mean (SD) | 2.98 (6.96) | | 0.08 (0.41) | | 0.93 (2.00) | 9.17 (10.84) | <0.01 |
| Gingival bleeding (% sites), mean (SD) | 36.08 (23.43) | | 30.09 (18.99) | | 34.43 (22.78) | 44.18 (25.80) | <0.01 |
| Plaque index (% sites), mean (SD) | 57.61 (28.14) | | 46.43 (27.53) | | 56.25 (27.23) | 69.49 (25.76) | <0.01 |
| Calculus index, mean (SD) | 0.91 (0.70) | | 0.52 (0.53) | | 0.86 (0.66) | 1.34 (0.66) | <0.01 |

Differences across groups determined by ANOVA (continuous) or χ^2^ (categorical) tests. P-values were adjusted for multiple comparisons using the Bonferroni correction.
