## Supplementary Table S2 for "A population-scale landscape of the subgingival microbiome reveals divergent routes to periodontal dysbiosis"

**Supplementary** **Table S1.** Characteristics of participants by periodontal health status according to custom-defined categories.

|  | **Periodontal Health (Custom Categories)** | | | | | |
| --- | --- | --- | --- | --- | --- | --- |
|  | **Health**  **(n=220)** | **Gingivitis**  **(n=430)** | **Periodontitis - Low Mean PD (n=235)** | **Periodontitis - Medium Mean PD (n=233)** | **Periodontitis - High Mean PD (n=234)** | **Adjusted P-value** |
| **Demographics** |  | | | | | |
| Age (years), mean (SD) | 56.00 (10.13) | 55.58 (9.44) | 57.67  (9.27) | 55.23  (9.46) | 54.24  (9.07) | 0.09 |
| Male (%) | 45.00 | 46.28 | 62.13 | 61.37 | 76.92 | <0.01 |
| Education (years), mean (SD) | 13.95 (2.31) | 13.89 (2.33) | 13.64  (2.53) | 13.61  (2.24) | 13.27  (2.12) | 0.11 |
| Non-White (%) | 5.45 | 6.98 | 5.96 | 6.01 | 12.82 | 0.48 |
| BMI (kg/m2), mean (SD) | 27.37 (4.73) | 28.91 (5.67) | 28.44  (5.31) | 28.99  (5.38) | 29.10  (5.43) | 0.20 |
| **Lifestyle and behavioral factors** |  | | | | | |
| Current smoker (%) | 9.55 | 8.88 | 9.79 | 16.74 | 27.78 | <0.01 |
| Lifetime smoking (pack-years),  mean (SD) | 12.97 (20.56) | 10.59 (18.83) | 13.70  (16.38) | 17.27  (19.55) | 27.60  (22.15) | <0.01 |
| Lifetime alcohol consumption (oz),  mean (SD) | 5144.44  (7514.52) | 5707.42  (13253.78) | 7708.99 (11003.14) | 11515.72 (34967.25) | 13762.59 (25213.47) | <0.01 |
| Alcohol consumption during the 12-month period 12–24 months prior (oz), mean (SD) | 112.99 (231.67) | 104.03 (284.27) | 139.94 (299.96) | 151.47  (328.17) | 194.69 (416.97) | 0.43 |
| Annual dental visit, (%) | 86.12 | 79.13 | 82.03 | 70.80 | 59.15 | <0.01 |
| Tooth brushing, every day (%) | 97.21 | 94.61 | 95.37 | 96.41 | 88.73 | 0.08 |
| Flossing, every day (%) | 31.13 | 16.75 | 29.30 | 23.29 | 14.56 | <0.01 |
| Exercise hours per week, mean (SD) | 5.23 (1.66) | 5.19  (1.67) | 5.22  (1.81) | 4.97  (1.88) | 4.94  (1.93) | 1.00 |
| **Medical history and medication use** |  | | | | | |
| History of myocardial infarction (%) | 30.91 | 28.84 | 34.47 | 45.92 | 65.38 | <0.01 |
| Self-reported hypertension (%) | 31.96 | 33.88 | 33.76 | 40.52 | 41.63 | 1.00 |
| Self-reported diabetes (%) | 5.96 | 10.05 | 9.87 | 12.55 | 9.91 | 1.00 |
| Systolic blood pressure (SBP) (mmHg),  mean (SD) | 117.24 (14.67) | 118.56 (14.86) | 118.91  (15.12) | 117.83  (14.89) | 118.29  (14.85) | 1.00 |
| Diastolic blood pressure (DBP) (mmHg),  mean (SD) | 71.74 (9.27) | 72.29 (8.86) | 71.95  (9.32) | 72.21  (9.73) | 72.94  (9.38) | 1.00 |
| Hypertension medication (%) | 43.64 | 40.70 | 46.38 | 53.88 | 60.26 | <0.01 |
| Lipid statin medication (%) | 20.00 | 21.86 | 24.68 | 36.91 | 45.30 | <0.01 |
| Diabetes medication (%) | 5.00 | 7.91 | 7.23 | 10.73 | 6.84 | 1.00 |
| Steroid medication (%) | 2.73 | 1.63 | 1.70 | 1.29 | 0.85 | 1.00 |
| **Blood glucose and lipid measures** |  | | | | | |
| Fasting blood glucose (mg/dL),  mean (SD) | 102.18 (28.97) | 104.97  (32.84) | 101.62  (25.06) | 104.24  (26.36) | 106.27  (28.07) | 1.00 |
| Self-reported high cholesterol (%) | 40.91 | 43.33 | 41.99 | 52.61 | 59.03 | 0.01 |
| Total cholesterol (mg/dL), mean (SD) | 205.26  (41.40) | 204.67 (41.18) | 202.67  (42.95) | 196.19  (45.56) | 195.00  (43.48) | 0.02 |
| Triglyceride (mg/dL), mean (SD) | 134.48  (77.07) | 142.36 (92.82) | 146.82  (91.78) | 154.16  (88.02) | 160.13  (96.78) | 0.12 |
| **Periodontal clinical parameters** |  | | | | | |
| Number of lost teeth, mean (SD) | 4.93 (5.95) | 5.50  (6.20) | 3.99  (4.14) | 4.29  (4.51) | 5.61  (5.50) | 0.02 |
| Pocket depth (PD), mean (SD) |  | | | | | |
| Whole mouth mean (mm) | 1.83 (0.31) | 1.86 (0.41) | 1.89 (0.16) | 2.29 (0.11) | 3.11 (0.62) | <0.01 |
| Maximum (Deepest site) (mm) | 4.01 (0.78) | 4.11 (0.92) | 6.19 (1.36) | 6.42 (1.40) | 7.75 (1.64) | <0.01 |
| Clinical attachment loss (CAL),  mean (SD) |  | | | | | |
| Whole mouth mean (mm) | 2.07 (0.48) | 2.11 (0.71) | 2.21 (0.49) | 2.63 (0.61) | 3.83 (1.36) | <0.01 |
| Maximum (mm) | 4.99 (1.29) | 5.08 (1.53) | 6.70 (1.99) | 7.33 (2.13) | 9.28 (2.46) | <0.01 |
| Alveolar crestal height (ACH),  mean (SD) |  | | | | | |
| Whole mouth mean (mm) | 1.65 (0.53) | 1.52 (0.55) | 1.84 (0.61) | 1.97 (0.73) | 2.59 (0.98) | <0.01 |
| Maximum (mm) | 4.34 (1.53) | 4.27 (1.54) | 4.84 (1.74) | 5.24 (1.99) | 6.77 (2.63) | <0.01 |
| Number of sites with PD ≥ 5 mm,  mean (SD) | 0.09 (1.22) | 0.05 (0.51) | 2.19 (1.56) | 3.69 (2.75) | 15.28 (14.88) | <0.01 |
| % Sites with PD ≥ 5 mm, mean (SD) | 0.06 (0.78) | 0.05 (0.45) | 1.57 (1.08) | 2.70 (2.03) | 12.84 (12.31) | <0.01 |
| Gingival bleeding (% sites), mean (SD) | 10.48 (5.78) | 42.66 (16.97) | 33.42  (22.38) | 38.86  (23.62) | 48.00  (26.61) | <0.01 |
| Plaque index (% sites), mean (SD) | 37.65 (24.70) | 58.47 (26.78) | 53.72  (25.79) | 59.92  (25.91) | 76.21  (24.64) | <0.01 |
| Calculus index, mean (SD) | 0.48 (0.49) | 0.76 (0.66) | 0.76 (0.58) | 1.09 (0.64) | 1.57 (0.59) | <0.01 |

Custom-defined categories: Periodontitis was defined as ≥1 site with both PD ≥5 mm and CAL ≥3 mm. Among participants not meeting this definition, gingival bleeding <20% was classified as periodontal health and ≥20% as gingivitis. Participants with periodontitis were further stratified into low, medium, and high mean PD groups based on tertiles of whole-mouth mean PD. Differences across groups were determined by ANOVA for continuous variables and χ² tests for categorical variables. P-values were adjusted for multiple comparisons using the Bonferroni correction.
