## Supplementary Table S3 for "A population-scale landscape of the subgingival microbiome reveals divergent routes to periodontal dysbiosis"

**Supplemental** **Table S2.** Characteristics of the Participants by Periodontal Health Status According to 2017 AAP Staging

|  | **AAP Staging** | | |
| --- | --- | --- | --- |
|  | **Stage I/II**  **(n=475)** | **Stage III/IV**  **(n=880)** | **Adjusted P-value** |
| **Demographics** |  | | |
| Age (years), mean (SD) | 53.82 (9.33) | 56.77 (9.45) | <0.01 |
| Male (%) | 41.68 | 64.66 | <0.01 |
| Education (years), mean (SD) | 14.07 (2.21) | 13.49 (2.36) | <0.01 |
| Non-White (%) | 5.26 | 8.64 | 0.92 |
| BMI (kg/m2), mean (SD) | 28.15 (5.38) | 28.89 (5.39) | 0.62 |
| **Lifestyle and behavioral factors** |  | | |
| Current smoker (%) | 8.03 | 16.93 | <0.01 |
| Lifetime smoking (pack-years), mean (SD) | 8.86 (17.10) | 19.22 (20.98) | <0.01 |
| Lifetime alcohol consumption (oz), mean (SD) | 5147.67 (10996.07) | 10050.30 (23811.84) | <0.01 |
| Alcohol consumption during the 12-month period 12–24 months prior (oz), mean (SD) | 106.77 (279.87) | 150.67 (331.66) | 0.57 |
| Annual dental visit, (%) | 83.52 | 71.84 | <0.01 |
| Tooth brushing, every day (%) | 96.04 | 93.69 | 1.00 |
| Flossing, every day (%) | 20.09 | 23.25 | 1.00 |
| Exercise hours per week, mean (SD) | 5.18 (1.69) | 5.09 (1.83) | 1.00 |
| **Medical history and medication use** |  | | |
| History of myocardial infarction (%) | 25.47 | 46.93 | <0.01 |
| Self-reported hypertension (%) | 31.29 | 38.70 | 0.26 |
| Self-reported diabetes (%) | 6.99 | 11.34 | 0.40 |
| Systolic blood pressure (SBP) (mmHg), mean (SD) | 117.52 (14.24) | 118.69 (15.26) | 1.00 |
| Diastolic blood pressure (DBP) (mmHg), mean (SD) | 72.15 (9.09) | 72.31 (9.33) | 1.00 |
| Hypertension medication (%) | 37.68 | 53.47 | <0.01 |
| Lipid statin medication (%) | 17.89 | 34.55 | <0.01 |
| Diabetes medication (%) | 5.26 | 8.98 | 0.54 |
| Steroid medication (%) | 1.89 | 1.48 | 1.00 |
| **Blood glucose and lipid measures** |  | | |
| Fasting blood glucose (mg/dL), mean (SD) | 101.21 (26.37) | 105.62 (30.34) | 0.33 |
| Self-reported high cholesterol (%) | 38.48 | 51.68 | <0.01 |
| Total cholesterol (mg/dL), mean (SD) | 207.09 (39.87) | 198.46 (44.46) | 0.02 |
| Triglyceride (mg/dL), mean (SD) | 138.68 (89.97) | 151.62 (90.65) | 0.70 |
| **Periodontal clinical parameters** |  | | |
| Number of lost teeth, mean (SD) | 3.88 (5.26) | 5.55 (5.52) | <0.01 |
| Pocket depth (PD), mean (SD) |  | | |
| Whole mouth mean (mm) | 1.84 (0.39) | 2.31 (0.63) | <0.01 |
| Maximum (Deepest site) (mm) | 4.04 (0.68) | 6.25 (1.88) | <0.01 |
| Clinical attachment loss (CAL), mean (SD) |  | | |
| Whole mouth mean (mm) | 1.88 (0.34) | 2.85 (1.08) | <0.01 |
| Maximum (mm) | 4.37 (0.76) | 7.66 (2.28) | <0.01 |
| Alveolar crestal height (ACH), mean (SD) |  | | |
| Whole mouth mean (mm) | 1.50 (0.48) | 2.06 (0.84) | <0.01 |
| Maximum (mm) | 3.71 (1.02) | 5.68 (2.19) | <0.01 |
| Number of sites with PD ≥ 5 mm, mean (SD) | 0.10 (0.37) | 5.62 (9.86) | <0.01 |
| % Sites with PD ≥ 5 mm, mean (SD) | 0.06 (0.24) | 4.55 (8.22) | <0.01 |
| Gingival bleeding (% sites), mean (SD) | 30.62 (19.87) | 39.03 (24.66) | <0.01 |
| Plaque index (% sites), mean (SD) | 47.65 (26.56) | 62.97 (27.49) | <0.01 |
| Calculus index, mean (SD) | 0.57 (0.55) | 1.10 (0.70) | <0.01 |

Differences across groups determined by Student t-test (continuous) or χ^2^ (categorical) tests. P-values were adjusted for multiple comparisons using the Bonferroni correction.
